# Diauxie and co-utilization are not exclusive during growth in nutritionally complex environments

**DOI:** 10.1101/752832

**Authors:** Elena Perrin, Michele Giovannini, Francesca Di Patti, Barbara Cardazzo, Lisa Carraro, Camilla Fagorzi, Veronica Ghini, Paola Turano, Renato Fani, Marco Fondi

## Abstract

The classic view of microbial growth strategy when multiple carbon sources are available states that they either metabolize them sequentially (diauxic growth) or simultaneously (co-utilization). This perspective is biased by the fact that this process has been mainly analysed in over-simplified laboratory settings, i.e. using a few model microorganisms and growth media containing only two alternative compounds. Models concerning the mechanisms and the dynamics regulating nutrients assimilation strategies in conditions that are closer to the ones found in natural settings (i.e. with many alternative carbon/energy sources) are missing. Here, we show that bacterial co-utilization and sequential uptake of multiple substrates can coexist when multiple possible nutrients are provided in the same growth experiment, leading to an efficient exploitation of nutritionally complex settings. The order of nutrient uptake is determined by the actual biomass yield (and growth rate) that can be achieved when the same compounds are provided as single carbon sources. Finally, using two alternative theoretical models we show that this complex metabolic phenotype can be explained by a tight regulation process that allows microbes to actively modulate the different assimilatory pathways involved.

## Introduction

Microorganisms must quickly and efficiently adapt to a variety of possible fluctuations in the surrounding environment. When considering changes in the pool of available nutrients, this is usually achieved by a tight regulation of their metabolic phenotypes by sensing the availability of specific compounds, synthesizing the enzymes required for their catabolism and repressing them after specific metabolites are depleted ^1^. The spectrum of possible bacterial metabolic adaptation strategies can be observed, for example, when growing cells in a medium containing a simple mixture of carbon sources. In this situation bacterial may exhibit different patterns including diauxic growth ^2^, simultaneous consumption ^3^, and bistable growth ^4,5^. Further, nutrients concentration and growth medium composition are known to affect other important cellular features such as motility ^6,7^ cell adhesion ^6^ and biofilm formation ^8,9^. Typically, these phenomena have been studied (both theoretically and experimentally) in model organisms (e.g. *Escherichia coli, Lactococcus lactis*) ^10^, grown on defined media containing simple mixtures of 2/3 carbohydrates (e.g. glucose and lactose) ^11–14^. In natural conditions, however, bacteria rarely encounter simple combinations of exploitable carbon/energy sources. Rather, complex mixtures of nutrients are common and often colonized by actively growing bacteria. Do the same models developed for simplified conditions hold also in real-case scenarios? At present, we witness a knowledge gap concerning the study of these processes in experimental settings that do not involve model organisms and/or defined media and we lack a sound theoretical understanding of the mechanisms driving nutrients assimilation strategies in conditions that are closer to the ones found in natural settings.

Bacterial exploitation of nutrient patches is made up of (at least) two different stages, i.e. physical interaction followed by carbon sources metabolic degradation. The capability of bacteria to interact with transient nutrient sources is well documented and has revealed their high efficiency in exploiting transient nutrient patches ^15,16^. Little is known, instead, on the molecular aspects regulating and influencing bacterial productivity once microscale nutrient hot spots are colonized. At this stage, i.e. when cells start to feed on the available carbon source(s), other cellular mechanisms need to be involved to ensure a systematic exploitation of the resource. Indeed, as nutrient patches are likely composed of complex nutrient mixtures (that may include carbohydrates, amino acids, lipids and nucleic acids) bacteria need to dynamically activate specific degradation pathways according to the kind and concentration of external nutrients. In other words, a continuous and flexible genetic reprogramming needs to be active to ensure that the preferred compound(s) are sequentially or simultaneously up-taken from the external environment and properly metabolized. Up to now, this latter aspect has been mostly overlooked despite it might be central in the understanding of micro-scale nutrients dynamics.

In this regard, the marine environment represents a paradigmatic example of the challenges encountered by microorganisms when it comes to the efficient (and rapid) exploitation of complex nutritional inputs. Such a habitat is thought to be characterized by a low average nutrient level (e.g., the concentration of amino acids is in the range of ∼10^−9^ M) and nutrients in general appear and disappear in a sporadic fashion, demanding a precise chemical response, a fast swimming speed, and ability to localize and exploit a nutrient patch once it is found ^6^. These are the conditions that are commonly faced by marine heterotrophic bacteria, i.e. those microorganisms relying on the assimilation of external biomass for both energy generation and nutrition. Their metabolism is pivotal for the maintenance and the correct balance of oceanic biogeochemical cycles as they are central to the so-called microbial loop, i.e. the trophic pathway of the marine food web responsible for the microbial assimilation of dissolved organic matter (DOM), by transforming phytoplankton-derived organic matter and fuelling the entire ocean biogeochemical nutrient cycle.

Here we have investigated the global regulation of a marine heterotrophic bacterium when grown in both a complex and a defined rich medium (i.e. including multiple possible carbon sources) using and integrating a set of complementary-omics techniques (i.e. transcriptomics and ^1^H-NMR metabolomics) with measured growth parameters. We show that the two main nutritional strategies commonly observed (co-utilization and sequential uptake of multiple substrates) can coexist in the same growth experiment, leading to an efficient exploitation of the available carbon sources. We also developed two theoretical models accounting for nutrients switching in a nutritionally rich environment in presence and absence of cell regulation acting at the level of resource allocation in the synthesis of nutrient assimilation pathways. We show that a model taking into consideration an overall regulatory control on the sequence of nutrients uptake produces a better fit with available experimental data in respect to a purely Michaelis-Menten kinetic model.

## Results

### Global regulation of a triauxic growth

*Pseudoalteromonas haloplanktis* TAC125 (hereinafter PhTAC125) cells were grown in shaken flasks in a complex medium composed of Schatz salts ^17^ and peptone as their C source. Optical density (OD) was measured every hour and cellular RNA was sampled in five different time points of their growth (Figure 1A). The growth curve displays a triauxic pattern (Figure 1A). An initial growth phase (growth rate of 0.023 h^-1^) is interrupted by a lag phase between min. 180 and min. 240; afterwards, cells start growing over but such growth is interrupted by another lag phase between min. 280 and min. 340. Cells then started growing again until the end of the experiment (growth rate of 0.026 h^-1^). The average growth rate across all the time points was estimated to be 0.01 h^-1^. To identify transcriptional changes during cell growth total RNA was extracted and sequenced (two biological and two technical replicates for a total of 20 samples) using Illumina MiSeq (Genomix4Life, Naples, Italy). The main features of the 20 sequenced samples are reported in Supplementary Material S1, Table S1. We clustered the genes according to their expression during the growth and identified 6 major trends (clusters C1 to C6, Figure 1B). Overall, we were able to cluster 2045 genes out of the 3448 encoded by the PhTAC125 genome (roughly 60%). We then performed a functional annotation and a functional enrichment analysis for the genes embedded in each cluster. One of these clusters (C6) did not include any significantly enriched functional category and thus it was discarded. Cluster C1 includes genes that display a decrease in their expression between the first two time points (T1 and T2) and a constant (low) expression across the rest of the growth curve.

**Figure 1:**
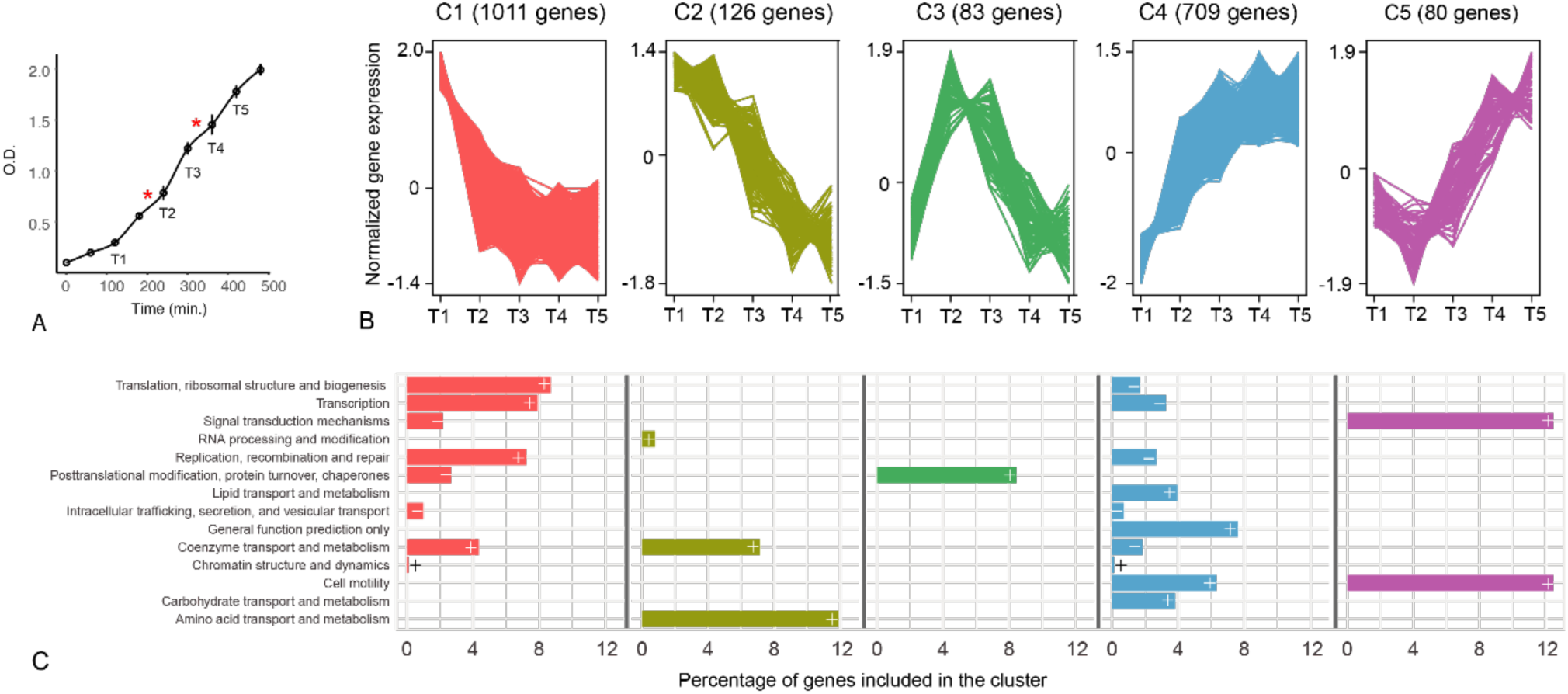
**A)** PhTAC125 growth curves in peptone. Asterisks represent hypothetical switch points and T1 to T5 indicated the sampled time points. **B)** Clustering of PhTAC125 genes according to their expression values (normalized according to *clust* algorithm, see Material and Methods) throughout the growth curve. **C)** COG functional categories that are significantly over- (“+” sign) or under- (“-” sign) represented inside each cluster.

Over-represented genes embedded in this cluster included those involved in basic housekeeping functions such as translation, DNA replication and transcription (Figure 1C). Genes embedded in Cluster 2 displayed a decreasing trend throughout the growth curve and mainly included genes involved in RNA processing, metabolism of coenzymes and amino acids transport and metabolism. The expression of the 83 genes included in Cluster 3 was characterized by an abrupt increase between T1 and T2 and then an overall decrease until the end of the curve. This cluster significantly included genes involved in post-translational modification, protein turnover and chaperons. Clusters 4 and 5 included genes whose expression tends to increase in the later stages of the growth; overrepresented genes in C4 mainly belong to lipid metabolism, cell motility and amino acids transport and metabolism. The expression of genes included in C5 decreases during the first stages of the growth and is then increased for the rest part of the curve. The cluster of genes includes those involved in signal transduction mechanisms and cell motility.

Whole-genome transcriptomics data depict a scenario in which PhTAC125 is active and fast-growing mainly during the first stages of the curve, as reflected by the relatively high expression of translation, transcription replication and coenzyme metabolism genes. Genes embedded in these categories are underrepresented among those increasing their expression in the last stages of the growth (Figure 1C) and over-represented among those with high expression values in the first stages of the growth. Metabolically, PhTAC125 cells seem to rely more on amino acids metabolism in the initial stages of their growth, consistently with their progressive exhaustion in the medium. The last part of the growth experiment was also characterized by an increase in gene expression of cell motility-related genes (overrepresented in C4 and C5). Finally, genes generally related to post-processing mechanisms peak their expression at T2.

### A non-*E. coli*-like regulatory response to nutrients exhaustion

The triauxic growth curve reported in Figure 1A suggests the presence of a dynamic control on the adjustment of cell physiology. Here we sought to quantify the regulatory effort required to growing cells for modulating such cellular response. We focused on transcriptional factors (TFs) and two component response systems (TCRSs) and analysed differentially expressed genes among three points of PhTAC125 growth curve, namely T1 vs. T3 and T3 vs. T5. These points should capture PhTAC125 cells during exponential growth after the first growth lag (T1), in-between the two growth lags (T3) and after the final growth lag but before getting to plateau (T5).

First, we checked whether PhTAC125 regulation system somehow resembled the model scheme of the known overall metabolic regulation (i.e. the one characterized in *E. coli*). Of the 81 transcription factors known to directly or indirectly control central metabolic enzymes ^18^, we found a reliable homolog (E-value<1e^-20^) only for 34 of them (Supplementary Material S1, Table S2). PhTAC125, for example, lacks key players in bacterial diauxic shifts as the major global regulator of catabolite-sensitive operons (when complexed to cAMP) *crp* and the genes responsible for the synthesis of cyclic AMP (adenylate cyclase, *cyaA*). Among the 34 global regulators identified, only 10 (roughly 25% of the shared ones and 12% of the entire *E. coli* set) displayed a significantly altered expression following the first transition (T1-T3) and none of them was differentially expressed following the second one (T3-T5). Details on the shared, differentially expressed TFs are provided in Supplementary Material S1 (Table S3).

A similar situation was observed for 8 selected sigma-factors that control gene expression globally ^18^. In this case, as expected, an ortholog was found for each of them, but only two of them showed an altered expression following T1-T3 transition and the expression of none of them was significantly altered following T3-T5 one. The two genes displaying a significant change in gene expression were *rpoS* and *rpoD*. RpoS is the primary regulator of stationary phase genes, whereas RpoD is the primary sigma factor during exponential growth. Expectedly, the first resulted to be up-regulated following the T1 to T3 transition whereas the second was down-regulated (Supplementary Material S1, Table S4).

With the exception of RpoS and RpoD, whose expression is in line with the global control of exponential vs. stationary phases, it appears that growth lags are regulated by mechanisms that poorly overlap with our current knowledge.

For this reason, we evaluated the expression of the entire repertoire of PhTAC125 TFs across the two points that involved the ceasing of cellular growth in our experiment, i.e. T1-T3 and T3-T5. Overall, we identified 41 differentially expressed TFs, 22 down-regulated and 19 up-regulated (Figure 2B and Supplementary Information S1, Figure S1 and Table S5) following the first growth interruption. The second growth lag was characterized by the significant change in expression (upregulation) of just one TF. Together with TFs, two-component regulatory systems (TCRSs) are a basic stimulus-response coupling mechanism to sense and react to changes in environmental conditions, e.g. nutrient concentration. We identified differentially expressed TCRSs in the two selected contrasts. Overall, we found 21 TCRSs-related genes that were differentially expressed in T1 vs. T3 and none in the T3 vs. T5 transition (Figure 2C, and Supplementary Information S1, Table S6).

**Figure 2:**
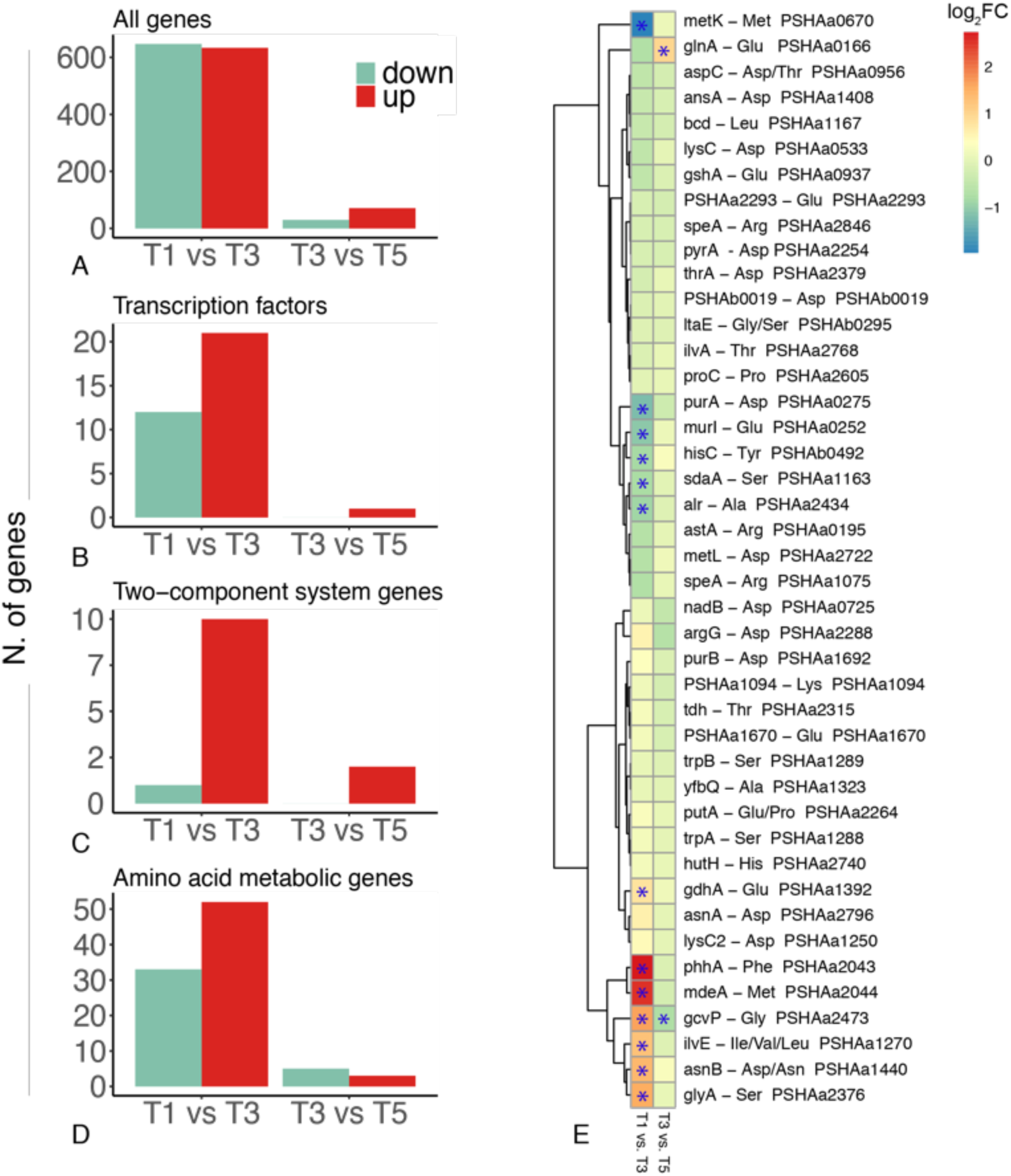
**A-D)** Number of up- and down-regulated genes in the two contrasts considered, for different gene categories **E)** Fold change of each gene responsible for the first degradative step of each amino acid. Differentially expressed genes are marked with an asterisks.

We conclude that the two growth lags observed in the curve apparently point to different reprogramming efforts that, in turn, may underpin distinct nutrients uptake strategies. The first growth interruption seems to have a deeper impact on the entire metabolic system, whereas the second could imply a fine tuning of the catabolic machinery. This is further confirmed by the overall number of DEGs across the selected contrasts (Figure 2A), 1280 between T1 and T3 (633 and 647 up- and down-regulated genes, respectively) and only 101 between T3 and T5.

### Preferential amino acid assimilation pathways and their (dis)regulation

Previous experiments have shown that PhTAC125 displays a coordinated sequence of amino acids degradation when grown in medium embedding complex mixtures of such molecules (i.e. peptone or casamino acid-based media) ^19^. In other words, some amino acids are preferred over others and are metabolized early in PhTAC125 growth curve. This switching among nutrients suggests that an active and modulated reprogramming occurs during PhTAC125 growth in a nutritionally complex environment. Similarly, differentially expressed amino acid metabolic genes (hereinafter AA-genes) are unevenly distributed among the two contrasts considered (Figure 2D). T1 vs T3 displays a higher number of DEGs (85, 33 down-regulated and 55 up-regulated) in respect to T3 vs T5 (8, 5 down-regulated and 3 up-regulated). Considering the amino acid assimilation pathways of differentially expressed AA-genes in the T1 vs T3 contrast (Figure S2), we didn’t observe a clear functional bias towards specific routes. Almost all the pathways are represented, both in terms of up- and down-regulated genes. Similarly, the switch between T3 and T5 included down-regulated genes involved in a broad spectrum of metabolic pathways including Val, Leu and Ile degradation, Tyr metabolism and Ala, Asp and Glu metabolism (one gene for each pathway) and up-regulated genes in Gly, Ser and Thr metabolism (1 gene), Lys biosynthesis (1 gene) and Arg biosynthesis (3 genes).

To unravel the faith of each amino acid inside the cell, we analysed the expression of the genes involved in all their possible first assimilatory step (Figure 2E and Supplementary Material S1, Table S7). We found 13 DEGs in the T1-T3 contrasts and 2 DEGs in the T3-T5 contrast. The first set comprised 7 up-regulated and 6 down-regulated genes; up-regulated genes were involved in Glu, Phe, Met, Gly Ile/Val/Leu, Asn/Asp and Ser degradation, whereas down-regulated genes were responsible for the first assimilatory step of Met, Asp, Glu, Tyr, Ser and Ala. DEGs identified in the second contrast included genes involved in Glu and Gly degradation. Taking the DEGs indicated above as a proxy for the entire assimilatory process of the corresponding amino acids, we noticed a good overlap with available PhTAC125 physiological data ^19^.

DEGs analysis also allowed the identification of the major amino acid entry points into PhTAC125 metabolism. We counted, for example, twelve alternative possibilities steps to metabolize Asp in PhTAC125 (Figure 2E), but the expression of only two of them (*purA* and *asnB*) appeared to be significantly modified during PhTAC125 growth. Similarly, 6 alternative steps can convert Glu to other cellular intermediates following its uptake. At T1, only two of these genes show an altered expression level, suggesting that these may represent the most relevant players in Glu assimilation and usage. Nearly the same holds for Ser, with 5 distinct entry points and only two of them being differentially regulated.

Overall, we have identified possible key players both in the switch among the set of metabolized amino acids and in the entrance of amino acids into PhTAC125 entire metabolic network. However, nutrients switching requires an efficient genetic regulation to ensure that each catabolic pathway is active at the right moment, allowing a correct proteome allocation. For this reason, we analysed the co-expression of genes belonging to the same metabolic pathway and identified an overall dis-regulation of such genes (average Fisher’s Z transformation average of Pearson correlation coefficient 0.49, Figure 3A,B and Figure S3, Table S8). Focusing on the known regulons including AA-genes (Table S9), we noticed that nearly half of them (3 out of 7) displayed a relatively low (0.47, ArgR) or almost absent (0.26 and 0.11, MetJ and TyrR1, respectively) correlation among the expression values of the corresponding genes. Figure 3C-E summarizes the details of the correlation existing among each gene of each pathway. In the case of ArgR regulon, for example, the major contribution to the low intra-regulon correlation is due to *astA* and *astD* (PSHAa0195 and 0196, respectively), showing an almost opposite expression pattern compared to the other ArgR regulated genes, especially with PSHAa2287-91 (Figure 3C). *astA* and *astD* are involved in the conversion of Arg to Glu, whereas ArgHA, B, C, F, G (encoded by PSHAa2287-91, respectively) are involved in the synthesis of Arg from Glu through the formation of citrulline and fumarate.

**Figure 3.**
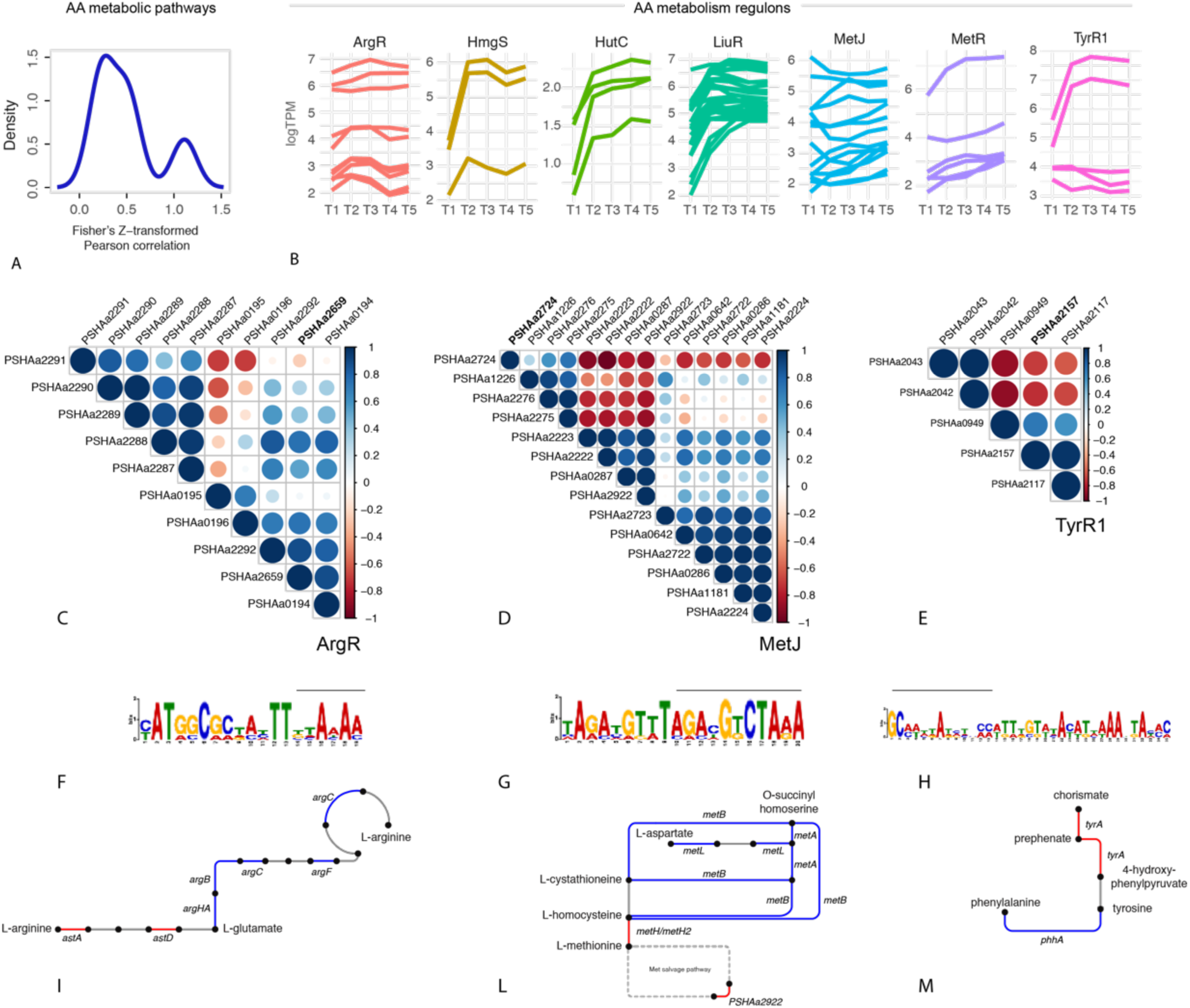
**A)** Cumulative density plot of Fisher’s Z transformation average of Pearson correlation expressing the co-regulation of genes belonging to the same pathway. B)Log2-transformed TPM values of the genes belonging to the same regulon. **C-E)** Graphic representation of the Pearson correlation matrix for the genes belonging to the regulons showing the lowest average correlation, Arg metabolism/biosynthesis, Methionine biosynthesis, Aromatic amino acids metabolism. Locus tags in bold indicate those of the regulator of the corresponding regulon. **F-H)** show the conserved upstream motif found for each gene of the considered regulon. The bar above each conserved motif indicates the overlap existing with the known TF binding site according to the RegPrecise database. Finally, in **I-M**) the reactions encoded by the genes included in the same regulon are schematically represented.

Concerning MetJ regulon, we noticed a group of genes (including PSHAa2222, PSHAa2223, PSHAa0287 and PSHAa2292) whose expression values are negatively correlated with those of genes PSHAa2274-76 and PSHAa1226 (Figure 3D). This group of co-regulated genes include those involved in the conversion of homocysteine to Met (PSHAa2222 and PSHAa2223), an L-alanine-DL-glutamate epimerase (PSHAa0287) and a Methylthioribulose-1-phosphate dehydratase involved in the Met salvage pathway (PSHAa2292). Finally, as for TyrR1regulon, PSHAa2042-43, coding for 4a-hydroxytetrahydrobiopterin dehydratase and phenylalanine-4-hydroxylase are negatively correlated to the other genes in the same regulon (Figure 3E). PSHAa2043 encodes *phhA* the gene responsible for the synthesis of Tyr from Phe, whereas PSHAa2042 (*phhB*) encodes a Pterin-4-alpha-carbinolamine dehydratase responsible for the conversion of 4a-hydroxytetrahydrobiopterin to dihydrobiopterin.

Upstream of most of the genes belonging to the three regulons considered, we were able to identify a conserved motif for each regulon (Figure 3F-H), partially overlapping with their known TF binding site. Finally, a closer inspection to the metabolic steps encoded by the differentially regulated genes of these regulons revealed that they usually belong to different and symmetric regions of the same metabolic pathway. PSHAa0195 and PSHAa0196 respectively encodes for *astA* and *astB*, responsible for the first steps of the route leading to the formation of Glu from Arg. The other genes of the ArgR regulon are mostly involved in the production of Arg starting from Glu (Figure 3I). Similarly, *phhA* (encoded by PSHAa2043) is involved in the formation of Tyr (from Phe), whereas all the other genes are responsible for the formation of Tyr from a set of different precursors (e.g. prephenate) (Figure 3M). In the case of MetR regulon, among the genes that could be reliably assigned to the methionine metabolic pathway, one of the two group of coregulated genes belong to the upper part of the pathway (upstream the main product methionine), whereas members the other one are in its close proximity (PSHAa2222) or involved in the methionine salvage pathway (PSHAa2292), the set of reactions responsible for the recycle of the thiomethyl group of S-adenosylmethionine from methylthioadenosine (Figure 3L).

Taken together these results suggest that i) the two growth lags observed (Figure 1A) may be the same phenotypic representation of two different cellular states (i.e. assimilation strategies) and that ii) a rather complex genetic regulation is at work to ensure a correct decision-making process in nutritionally dynamic environments. In the next sections these two aspects will be elucidated using controlled growth conditions and a combination of NMR and theoretical modelling.

### Combination of simultaneous and sequential amino acids uptake

Up to now, we have analysed the behaviour of bacterial cells in a complex medium, using gene expression as a proxy for amino acids assimilation pathways. The medium used (peptone) is a complex mixture of nutrients whose exact composition is unknown.

Accordingly, it is not possible to conclude that the observed growth features (i.e. triauxic growth) are due to the exhaustion of certain preferred amino acids in the medium. For this reason, we assembled a medium including 19 amino acids (named 19 AA medium, cysteine was not included in the list because of difficulties in its unambiguous quantitation during the experiments due to its spontaneous oxidation, as also reported in ^20^) and determined the kinetics of their usage during PhTAC125 growth. Data obtained revealed that an important fraction of all the provided amino acids (16 out of 19) are consumed in the first 7 hours of the growth. Afterwards, the remaining three amino acids (His, Met and Trp) are (slowly) metabolized (Figure 4A). Clustering the amino acids assimilation profiles allowed a clearer visualization of the order in which amino acids are used by PhTAC125 during its growth (Figure 4B). This analysis divided the set of metabolized compounds into four main, non-overlapping clusters. Gln, Glu and Arg are the first amino acids to be consumed in the medium. Their concentration reaches (negligible) values close to 0.01 mM after 4.5 hours of growth, remaining constant afterwards. The second set of amino acids is composed of Asn, Asp, Leu and Pro. Their degradation starts with a small delay in respect to the one of the first cluster, and they are completely removed from the medium only between 6 and 6,5 hours. The third cluster includes 9 amino acids (Figure 4B). Their consumption is rather slow in the first 3 hours of growth; afterwards, it accelerates leading to negligible concentration of the corresponding amino acids at 7.5 hours. The concentration of amino acids belonging to the fourth cluster remains overall constant for the first six hours of growth. After that moment, corresponding to the point in which all the other amino acids are consumed, it starts decreasing. Importantly, this pattern of amino acids assimilation results in a triauxic growth curve (Figure 4C). Indeed, (short) growth lag phases are observed after 4 and 6 hours of growth, in correspondence with the major transition in amino acids assimilation pattern. Overall, this behaviour highlights a balanced mix between simultaneous and sequential uptake of nutrients. Amino acids belonging to the same group (Figure 4D) are simultaneously metabolized by the cells but the assimilation of different groups occurs with different dynamics and is responsible for growth lags in the curve. Finally, a typical diauxic nutrient shift is observed when all the main (preferred) sources are exhausted and the degradation of the other (previously ignored) compounds begins. The order in which nutrients are used by microbes during the growth depends both on the final biomass and the specific growth rate achievable when grown with amino acids as sole carbon sources. Using available growth phenotypes ^21,22^. We investigated the correlation between amino acids consumption and the achievable biomass amount when the same carbon sources of the 19 AA experiment were used as single carbon and energy sources. Figure 4E shows that, on average, amino acids included in cluster 1 permit a higher cell density than those in cluster 2 and so on. Similarly, when focusing on single amino acids and the growth rate measured when using them as unique C source, higher growth rates are obtained with those amino acids that are degraded first in our 19 amino acids growth curve (Figure 4F). These data confirm that, when faced with multiple alternatives, bacterial metabolism has evolved in order to start feeding on those that ensure then most efficient growth, leaving the others for the latter stages of the growth ^23^.

**Figure 4.**
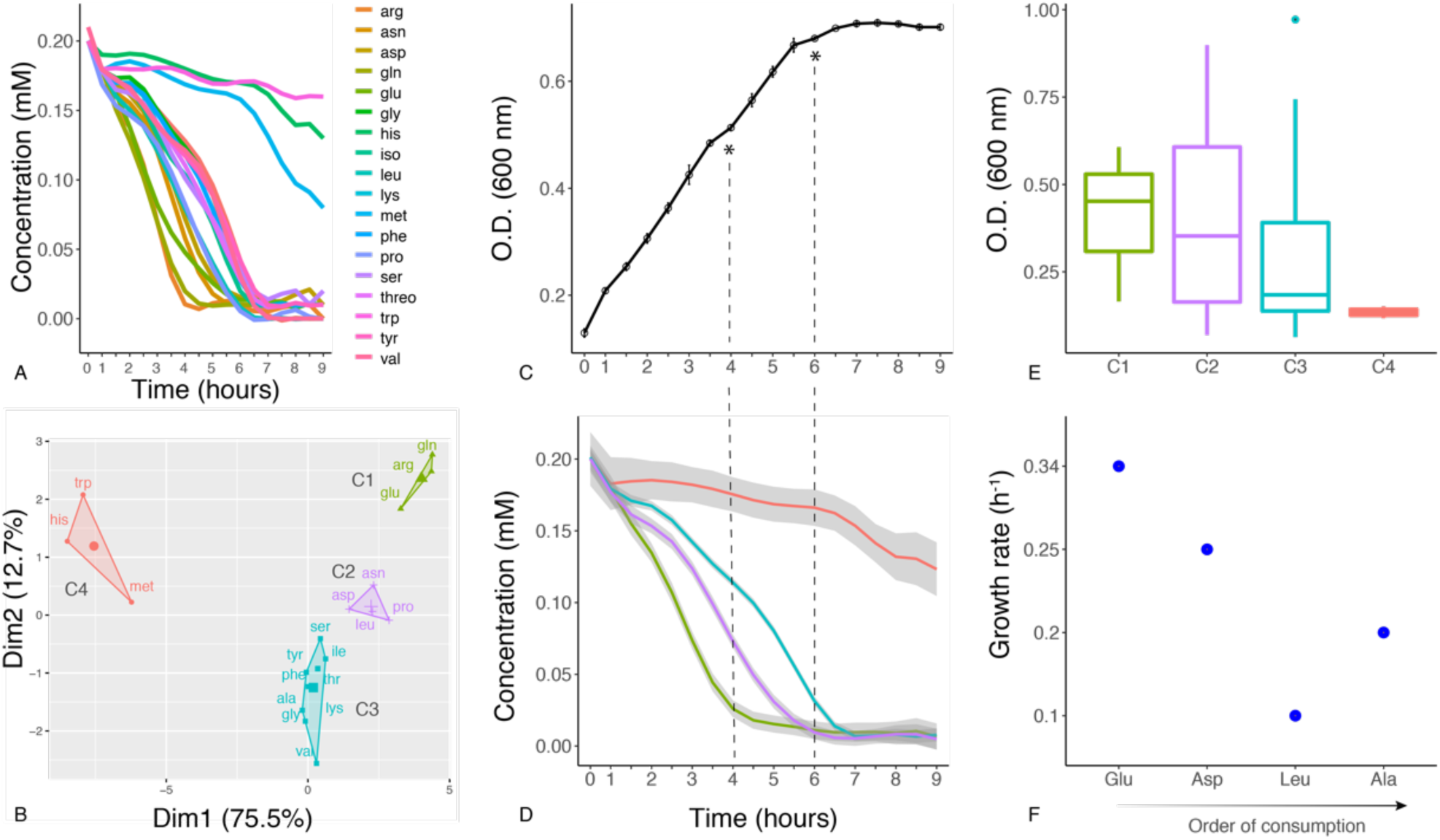
**A)** Degradation dynamics for each of the 19 amino acids included in the defined AA medium. **B)** Clustering of time-resolved concentration values for the 19 amino acids analysed. **C)** Growth curve of the 19 amino acids experiment. Asterisks indicate growth lags. **D)** Degradation dynamics for each of the 4 identified clusters of amino acids included in the defined AA medium. **E)** Average obtained biomass when using each of the amino acids of each cluster as sole carbon source. **F)** Growth rates obtained when using the amino acids indicated on the x-axis as sole carbon source.

At least two different explanations may account for the mixed sequential/diauxic nutrient uptake. Either this phenotype is “simply” determined by different uptake kinetics of the different compounds or the assimilation pattern is actively regulated by the cells. To discern between these two scenarios, we implemented two mathematical models accounting for cell growth and nutrients uptake during the 19 AA experiment. To reduce the complexity of the problem, the 19 amino acids were lumped into the 4 corresponding clusters shown in Figure 4B and D. In this way we modelled the growth of PhTAC125 in a hypothetical growth medium embedding four different groups of carbon sources ideally representing the 19 AA medium. The first model is based on the Michaelis-Menten Monod kinetics (MMM model) and is formulated as follows: 

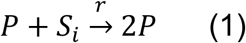

Where *P* represents bacterial cells, *S*_*i*_ (with *i* = 1, 2, 3, 4) represents each of the four groups of pooled C sources and *r* the rate at which the reaction occurs. Specifically, *r* was modelled according to a canonical (Michaelis-Menten-derived) Monod kinetics with: 

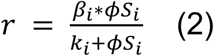

Where *β*_*i*_, *ϕS*_*i*_ and *k*_*i*_, represent the maximum rate constant for cell production, the concentration and the Michaelis-Menten constant for the *i*_*th*_ group of amino acids, respectively. According to these formulas, the state variables model can be written as: 

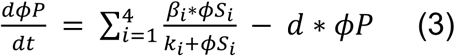

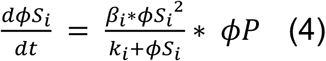

Where *S*_*i*_ (with *i* = {1, 2, 3, 4}) represents each of the four lumped substrates and *d* the bacterial cells death rate.

The second model implemented here accounts for the effect of the regulatory processes of catabolite inhibition and activation that can be observed during microbial growth on multiple substrates. The model is a modified version of the cybernetic model proposed by ^24^, overall resembling the one proposed in ^25^. According to this model, the assimilation of substrate *S*_*i*_ by cells *P* (equation (1)) is assumed to be catalysed by the set of enzymes *e*_*i*_ (with *i* = 1,2,3,4). The assumption here is that enzymes responsible for the assimilatory pathway of each pool of nutrients are induced by the presence of *S*_*i*_ (and repressed by the presence of the other nutrients). This alternative model can be written as: 

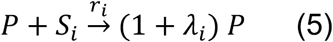

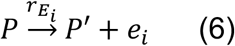

*P*′ represents the fraction of biomass produced minus the concentration of the key assimilatory enzyme (*e*_*i*_). To maintain consistency with equation (1), we assumed *P*′ to be negligible and that *λ*_*i*_ = 1. The rates *r*_*i*_ and 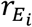 are expressed as follows: 

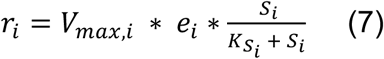

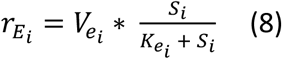

Where 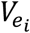 is the maximum rate constant for enzyme’s biosynthesis and *V*_*max,i*_ is the is the maximum rate constant for bacterial production *P* on the *i*-th substrate.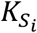 and 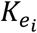 are the Michaelis-Menten constant for the *i*-th substrate and the synthesis of the *i*-th enzyme, respectively.

The inhibition/activation effect due to the concentration of the different substrates is accounted for by two (control) variables, *u*_*i*_ and *v*_*i*_, representing the fractional allocation of resource for the synthesis of *e*_*i*_ and the mechanism of controlling enzymes *e*_*i*_ activity, respectively. *u*_*i*_ is expressed as: 

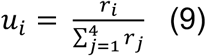

With 0 ≤ *u*_*i*_ ≤ 1 and 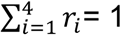. The other control parameter, *v*_*i*_ is expressed as: 

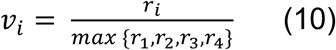

where the denominator accounts for the observation that priority is given to the consumption of the substrate(s) that guarantee the highest growth rate ^23^. The model further considers enzyme production rate (*β*_*i*_), the effect of dilution of the specific enzyme level due to cell growth (*α*), constant protein decay in the cells and bacterial death rate (*d*) and can be written as follows: 

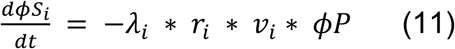

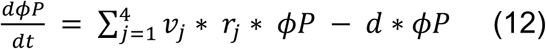

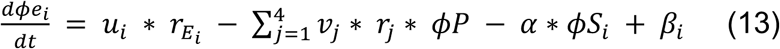

The model parameters were determined by fitting the experimental data (shown in Figure 4A-D) with model simulations and their values are reported in Supplementary Material S1 (Table S10 and S11). As shown in Figure 5, the cybernetic model accurately reproduces the dynamics of all the species considered. This is even clearer in the case of nutrient concentration dynamics where the model implementing the MMM model, is not capable of producing a satisfactory approximation of the real data. Indeed, R^2^ calculation indicates that the cybernetic modelling framework is performing better on 4 out of 5 of the species included in the model (Table 2).

**Table 1.** Fisher’s Z transformation average of Pearson correlation coefficient among the genes belonging to the same regulon

| *Regulon* | *Process* | *Fisher's Z transformation of Pearson correlation coefficient* |
| --- | --- | --- |
| *HutC* | His degradation | 2,55 |
| *HmgS* | Tyr degradation | 1,92 |
| *MetR* | Met biosynthesis | 1,22 |
| *LiuR* | Branched-chain amino acid degradation | 0,97 |
| *ArgR* | Arg biosynthesis/degradation | 0,47 |
| *MetJ* | Met metabolism, Met degradation | 0,26 |
| *TyrR1* | Aromatic amino acids metabolism | 0,11 |

**Table 2.** R^2^ calculation for the different simulations using the two different models implemented in this work.

| Model | Cluster A | Cluster B | Cluster C | Cluster D | Biomass |
| --- | --- | --- | --- | --- | --- |
| MMM model | 0.9844 | 0.9922 | 0.9584 | 0.9584 | 0.9959 |
| Cybernetic | 0.9920 | 0.9973 | 0.9953 | 0.9193 | 0.9967 |

**Figure 5:**
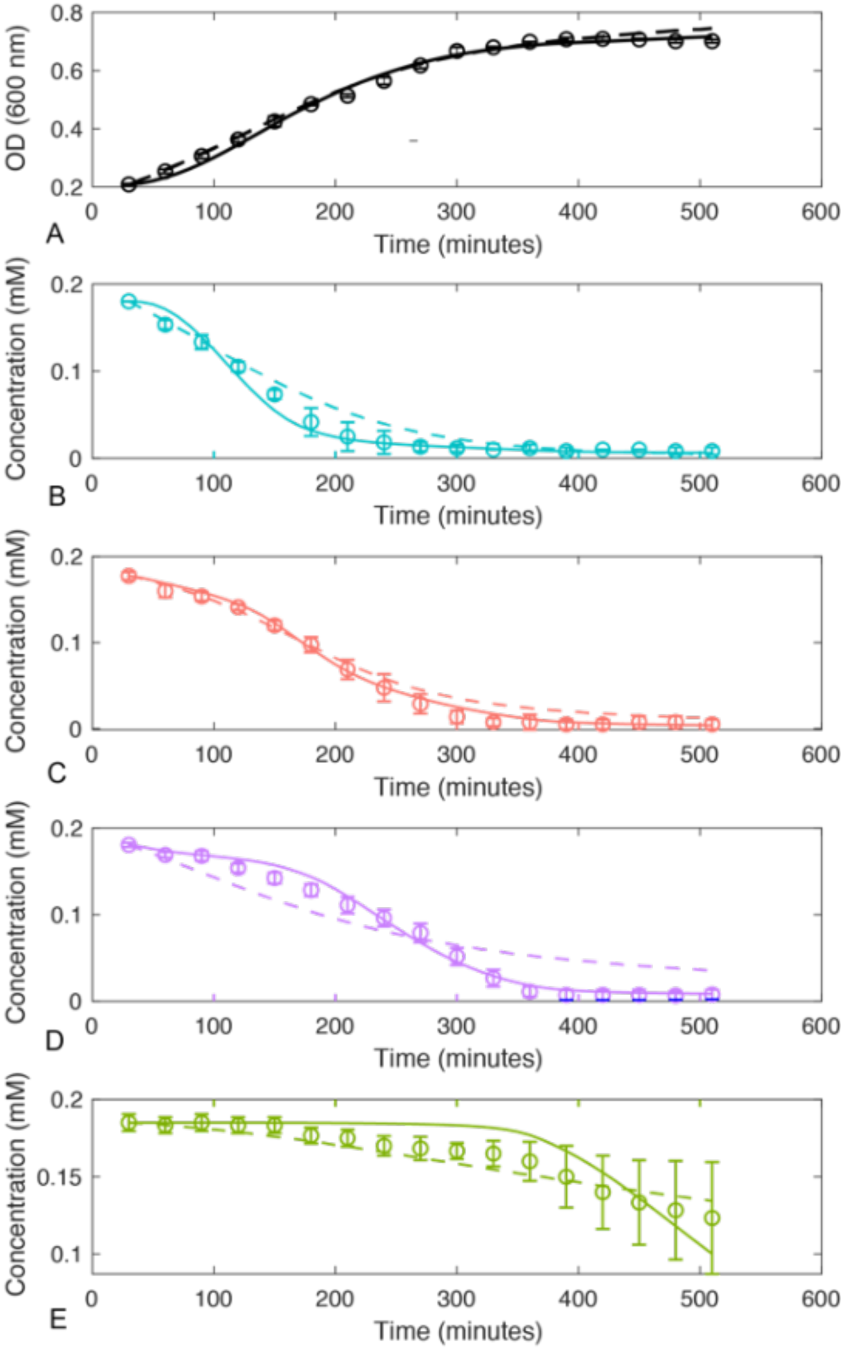
Simulation outcomes for the two models implemented here for biomass **(A)** and lumped nutrients **(B-E)**. Dashed lines represent the prediction for the MMM model. Solid lines represent the prediction for the cybernetic model. Circles represent experimental data (same data shown in Figure 4D).

These results suggest that, in the conditions tested, the uptake of nutrients is tightly regulated, leading to the simultaneous presence of diauxic and co-utilization strategies within the same growth curve. Hints on the main catabolic players involved in such assimilation patterns were obtained combining transcriptomics data from the complex-medium experiment and RT-PCR on specific targets (see Supplementary Material S1, Figure S4, Table S12).

## Discussion

Our knowledge on the possible bacterial strategies for nutrients assimilation when multiple sources are available is biased by the fact that it has been mainly studied in a few model organisms, providing them with a reduced number of possible inputs (compared to those available in their source environment). Here, we have characterized a non-model response to nutrients switching and studied the process of bacterial nutrients uptake in experimental conditions that more closely resemble a natural setting. Using a marine heterotrophic bacterium (*P. haloplanktis* TAC125) as a case study strain, we have shown that its response during growth lags do not resemble the one currently known for *E. coli*. Only 10% of the *E. coli* metabolic regulators and two (out of eight) main generic controllers (*rpoS* and *rpoD*) displayed an altered expression level in our experiments. The poor overlap between PhTAC1215 and *E. coli* transcriptional response suggests that, in marine bacteria, the response to nutritional switches and/or multi-auxic growth patterns may involve still untapped genetic circuits. As a matter of fact, PhTAC125 is known to lack the CCR system ^26^, which is currently referred to as the main driver in metabolic switches and diauxic phenotypes. Using time-resolved transcriptomics we have shown that growth lags in a nutritionally complex environment are probably due to the exhaustion of specific carbon sources and that such event has a deep impact also on other important gene categories including, for example, motility-related genes. In the second part of the curve, i.e. when nutrient concentration decreases, cell motility genes increase their expression, probably reflecting the need to explore the surrounding environment for other potential sources. This is in line with the observation that many bacteria become motile when nutrients are scarce ^27^. Moreover, among the genes peaking their expression in correspondence of the first growth lag, those involved in post-translational modification, protein turnover and chaperon are significantly over-represented. Generally speaking, proteins belonging to this functional category can be easily associated to the stress encountered by an exponentially-growing batch culture that exhaust (part of) the readily available and preferred nutrients in the growth medium. At this moment cells undergo a regulated transition into stasis by activating a stereotypic stress response. Post-translational modifications have been shown to play a role in the starvation-induced growth arrest, for example in *S. coelicolor* ^28^. It is to be noticed that these proteins do not increase their expression in the later stages of the growth and, in particular, in the second growth arrest experienced by PhTAC125. This, together with the observation that both the overall number of DEGs and of those involved in many other functional processes (amino acids degradation, TFs and TCRS) is much higher in correspondence of the first time contrast, suggested that the same phenotype (the two growth lags) were mirrored by profoundly different cellular reprogramming patterns. This prompted us to investigate more in depth the regulation of amino acid assimilation pathways. This analysis highlighted the tight regulation required to efficiently exploit complex and amino acid enriched, nutritional conditions. A paradigmatic example of this capability is the (dis)regulation of several genes belonging to the same regulon (ArgR, TyrR1 and MetR), responsible for the activity of distinct and conflicting functional metabolic modules inside the same metabolic pathway. In principle, all the genes belonging to the same regulon are under the control the same TF. It is known that some TFs may function as either activators or repressors, often according to the positioning of the TF binding site in the target promoter ^29,30^, although this feature had not been described, to date, for amino acid assimilation pathways. Apparently, such a mechanism is at work in some of the amino acid catabolic pathways of PhTAC125, probably ensuring an efficient and correct exploitation of the amino acid mixture available in the surrounding environment.

Despite being closer to the environmental natural setting, growth in complex medium does not allow a precise understanding of the usage of all the available carbon sources. For this reason, we have assembled a defined but nutritionally rich medium containing 19 amino acids as the sole carbon sources for the bacterial cells. Tracking their concentration in time, we showed that the two main feeding strategies commonly thought to be exclusive to each other (i.e. sequential and simultaneous uptake of nutrients) can co-exist in the same growth curve. Clustering metabolized amino acids into four major groups revealed that amino acids belonging to the same group are co-utilized, whereas the switch among the different clusters is tightly modulated. A “canonical” diauxic shift is finally observed at the end of the growth, when the consumption of a set of previously untapped nutrients begins. Thus, similar to the complex medium growth experiment (Figure 1), the two growth lags apparently underlay different cellular states. Indeed, despite the exhaustion of nutrients is common to both growth lags, in one case (the first lag) cells are already metabolizing alternative compounds when the exhaustion of the preferred sources occurs. The order in which nutrients are utilized can be explained by the biomass yield and growth rates obtained when each single amino acid is provided as single carbon sources. Those amino acids allowing the highest growth rate and biomass production are those that are consumed first in the 19 AA experiment. Finally, we have shown that the dynamics of nutrients degradation can be explained using a theoretical model that accounts for gene regulation and, in general, for the proper resource allocation for the synthesis of the main assimilatory pathways. This modelling framework can accurately interpret the pattern of nutrients degradation in a nutritionally rich environment.

In conclusion, we would like to stress the importance of cultivating and studying microorganisms in nutritional conditions that more closely resemble the ones most found in nature. By doing so and using a combination of computational and experimental (transcriptomics and NMR-based metabolomics) approaches we have shown that, despite diauxie and co-utilization strategies have been usually thought as conflicting phenotypes, they can co-exist in the same growth curve and give rise to a diversified ensemble of feeding strategies. It will be interesting to investigate which are the molecular mechanisms allowing the implementation of this mixed and apparently unconventional feeding strategy and, in particular, the fine-tuned regulatory circuits that are probably responsible for the efficient switching among all the available carbon sources. Future efforts will be also devoted to understanding the effect of fluctuations (in the number of cells and/or in nutrients concentration) and of possible population heterogenicity ^10,11^ on the resulting growth dynamics of heterotrophic marine bacteria.

## Material and methods

### Bacterial strain, media and growth condition

*Pseudoalteromonas haloplanktis* TAC125 ^31^ cells were routinely grown in Marine Agar (MA) or Broth (MB) (Condalab, Spain) under aerobic condition at 21°C. The stock suspension of the strain was stored in 20% [v/v] glycerol solution at -80°C. For growth curves experiment, Schatz salts ^32^ (1 g/l KH_2_PO4, 1 g/l NH_4_NO_3_, 10 g/l NaCl, 0.2 g/l MgSO_4_ ×7H2O, 0.01 g/l FeSO4×7H2O, 0.01 g/l CaCl2 ×2H2O) were supplemented with 5 g/l Peptone N-Z-Soy BL 7 (Sigma-Aldrich S.r.l) (complex medium) or with 19 amino acids (19 AA medium) each one at a final concentration of 0.2 mM (cysteine was not included due to its rapid oxidation to cystin ^20^). In both cases the pH was adjusted to 7.0. All the amino acids were purchased from Sigma-Aldrich S.r.l.. All the experiments were performed under aerobic condition at 21°C.

### Growth curve experiments

The growth curve experiments were performed after two precultures, as in Wilmes et al. ^19^ with some adaptations. For the complex medium growth curves, a first preculture was grown for 20 hours, in 20 ml MB medium in a 100 ml flask. Then this preculture was diluted 1:1000 in a final volume of 100 ml of the complex medium in a 1 l flask. After 20 hours of growth, the optical density (OD_600_) of this second preculture was measured to be used to inoculate the final flask (1l) in a final volume of 150 ml of Schatz salts and Peptone with a starting OD_600_ ∼0.1. For the growth curves in the 19 AA medium, the first preculture in MB was diluted 1:100 in a final volume of 100 ml of the 19 AA medium in a 1 l flask. After 22 hours of growth, the second preculture was washed, resuspend and used to inoculate the final flask (1l) in a final volume of 200 ml of the 19 AA medium with a starting OD_600_ ∼0.1. In both experiments, the pre-cultures and the final growth curves were incubated at 21°C with shaking. Each experiment was performed in duplicate. Cell growth was monitored measuring the OD_600_ every hour in the experiments in complex medium, and every half an hour in the experiments with the 19 AA medium. Three different measures were performed at each time point for each biological replicate.

### Sampling

Two biological replicates of the growth curves performed in complex medium were used for RNA-seq experiment. Every hour, in correspondence of the OD_600_ measurements, 2 replicates of 500 µl each for each curve, were treated with the RNA protect bacteria reagent (Qiagen S.r.l.) and conserved at -80°C.

During the experiment in the 19 AA medium, at each time point, 2 replicates of 500 µl each for each curve, were treated with the RNA protect bacteria reagent like above, while 2 replicates of 1 ml each for each curve were filtered (Filtropur 0.2 µm, SARSTED AG & Co. KG) to remove bacterial cells and conserved at -20°C for NMR metabolomic.

### RNA extraction and sequencing

For RNA-seq, a preliminary sequencing (data not shown) was performed on an Illumina Hiseq 50 platform (Genomix4Life S.r.l., Italy). Total RNA was extracted with a RNeasy Tissue Mini Kit (Qiagen S.r.l.) following manufacturer’s instructions. For improving the lysis step proteinase k and lysozyme were added to the lysis solution and the samples were homogenized using Tissue lyser II (Qiagen S.r.l.). The concentration and purity of RNA were analyzed using a NanoDrop ND-1000 (Thermo Fisher Scientific) and a Bioanalyser (Agilent Technologies, Inc.). rRNA was removed from the sample using the Ribo-Zero Magnetic Kit (Bacteria) (Illumina, Inc.). The quality of the RNA depletion was then checked using Bioanalyzer (Agilent RNA 6000 PICO Assay, Agilent Technologies, Inc.). The ScriptSeq v2 RNA-Seq Library Preparation Kit (Illumina, Inc.) is then used to make the RNA-Seq library from the Ribo-Zero treated RNA. For each library 1 µg of RNA (rRNA depleted) was used following manufacturer’s instructions. The quality of the libraries was evaluated using Bioanalyzer (Agilent Technologies, Inc.).

For the final experiment total RNA was extracted from a total of 20 samples of the growth curve in complex medium (5 time points, 2 technical and two biological replicates) and library sequencing has been carried out at Genomix4Life S.r.l. (Italy) on an Illumina NextSeq500 (single-end sequencing strategy, 1 × 75bp, ∼25 reads/sample).

For Real-Time PCR (qRT-PCR), total RNA was extracted from the samples of five time points (T4, T6, T8, T10 and T12) for each biological replicate of the growth curve performed in the 19 AA medium, using a RNeasy Mini Kit (Qiagen S.r.l.), following the manufacturer’s instruction. DNA was then removed from the samples using a RNase-free DNase (Qiagen S.r.l.). 10 µl of the extracted RNA was reverse-transcribed using a Superscript II Reverse Transcriptase (Invitrogen) with Random primers (Invitrogen) following the manufacturer’s instruction.

### Quantitative Real-Time PCR (qRT-PCR)

qRT-PCR reactions were performed in a final volume of 10 μl containing 1 μl of a 1:10 dilution of each cDNA, 5μl of Powrup Sybr Master Mix (Life Technologies) and 1 μM of each primer (Supplementary Material S1, Table S12). Primers were designed using the Primer3 software ^33^. Each sample was spotted in triplicate.

A first experiment using known amounts of DNA of the PhTAC125 strain (1-0.1-0.01-0.001 ng) were performed to obtain a standard curve and calculate the amplification efficiency for each primer pairs (data not shown). *rplM* and *dnaA* genes were used as internal references to normalize mRNA content.

All the reactions were performed on a QuantStudio™ 7 Flex Real-Time PCR System (Applied Biosystems by Life Technologies). Cycling conditions were: hold stage [50°C for 2’ and 95°C for 10’], PCR stage [40 cycles of: 95°C for 30’’, 59°C for 1’, 72°C for 15’’, melt curve stage [95°C for 15’’, 60°C for 1’, 95°C for 15’].

### RNAseq data analysis

Bowtie 2 (v2.2.3) ^34^ was used to align raw reads to *P. haloplanktis* TAC125 reference genome (GCA_000026085.1_ASM2608v1). rRNA depletion, strand specificity and gene coverage were evaluated using BEDTools (v2.20.1) ^35^ and SAMtools (v0.1.19) ^36^ to verify the library preparation and sequencing performances. Raw read counts were then used to calculate TPM values for each PhTAC125 gene. Clusters of co-regulated genes were identified using the Clust tool ^37^ using the following parameters: k-means clustering method, tightness weight equal to 0.3 and Q3s outliers threshold equal to 2.0.

Differentially expressed genes between the various contrasts were identified using the R (R Development Core Team, 2012, https://www.r-project.org/) package DeSeq2 ^38^ using default parameters and the following thresholds: adjusted p-value < 0.01 and log2FC > 0.75 or log2FC < -0.75. The clustering of genes based on their FC was performed using the Pheatmap R package. Visualization of Pearson correlation was performed using ‘corplot’ R package.

### Functional enrichment analysis and regulon identification

To conduct functional enrichment, each gene whose upstream intergenic region was clustered in one of the three clusters was assigned to a specific functional category using a BLAST ^39^ search against the COG database ^40^, with default parameters and considering a hit as significant if E-value < 1e-20. The exact binomial test implemented in the R package was used to assess over- and under-represented functional categories against the corresponding genomic background. Available information on PhTAC125 amino acid metabolism regulons were retrieved using RegPrecise database ^41,42^ The RegPrecise includes information for 7 PhTAC125 regulons (Table S9).

### Motif finding

Shared, conserved upstream motifs were searched up to 200 bp upstream of the genes belonging to the same regulon. These sequences were retrieved were retrieved from the *P. haloplanktis* TAC125 reference genome and fed into the MEME suite ^43^, version. MEME was used in combination with MAST(version 5.0.5) ^44^ for identifying the most plausible shared motifs upstream of the selected genes. MEME was used setting the following parameters: -nmotifs 5, -minw 6, -maxw 30, -objfun classic, -revcomp, -markov_order 0, -minsites 1, -maxsites 3. All the other parameters were set as default. MAST was used using default parameters. In all cases, only the best scoring motif was considered for further analyses, provided that the search produced a significant result (e-value < 0.05). The conservation of identified shared motifs was represented using WebLogo ^45^.

### Modelling

The deterministic system was simulated by numerically integrating differential equations using the Matlab built-in function ode45 v. 2019a. To estimate the unknown parameters of the model from experimental data we used a stochastic curve-fitting *in-house* Matlab software. The algorithm is based on the paper by Cardoso et al. ^46^ and consists in the combination of the non-linear simplex and the simulated annealing approach to minimize the squared deviation function. The codes used to perform the simulations reported in this work and the details about the options of the curve fitting environment, are available at https://multiauxic.sourceforge.io.

### Metabolomics assay and data analysis

^1^H-NMR spectra were acquired on cell media to monitor the uptake of the various amino acids by measuring their levels in samples collected at different time points of cell growth. Samples were prepared in 5.00 mm NMR tubes by mixing 60 μL of a potassium phosphate buffer (1.5 M K_2_HPO_4_, 100% (v/v) ^2^H_2_O, 10 mM sodium trimethylsilyl [2,2,3,3−^2^H_4_]propionate (TMSP), pH 7.4) and 540 μL of sample.

Spectral acquisition and processing were performed according to procedures developed at CERM ^47–52^. All the spectra were recorded using a Bruker 600 MHz spectrometer (Bruker BioSpin) operating at 600.13 MHz proton Larmor frequency and equipped with a 5 mm PATXI ^1^H-^13^C-^15^N and ^2^H-decoupling probe including a z axis gradient coil, an automatic tuning-matching (ATM) and an automatic and refrigerate sample changer (SampleJet). A BTO 2000 thermocouple served for temperature stabilization at the level of approximately 0.1 K at the sample. Before measurement, samples were kept for 5 minutes inside the NMR probe head, for temperature equilibration at 300 K.

^1^H-NMR spectra were acquired with water peak suppression and a standard NOESY pulse sequence using 128 scans, 65536 data points, a spectral width of 12019 Hz, an acquisition time of 2.7 s, a relaxation delay of 4 s and a mixing time of 0.1 s. The raw data were multiplied by a 0.3 Hz exponential line broadening before applying Fourier transformation. Transformed spectra were automatically corrected for phase and baseline distortions. All the spectra were then calibrated to the reference signal of TMSP at δ 0.00 ppm using TopSpin 3.5 (Bruker BioSpin srl). The signals deriving from each amino acid were assigned using an internal NMR spectral library of pure organic compounds. Matching between the present NMR spectra and the NMR spectral library was performed using the AMIX software. The relative concentrations of the various amino acids were calculated by integrating the corresponding signals in defined spectral range, using a home-made R 3.0.2 script.

## Funding

This project was supported by a PNRA (Programma Nazionale di Ricerca in Antartide) grant (grant PNRA16_00246).

## Authors contribution

MF, EP and RF initiated the project. MF and EP designed all the experiments. EP performed and/or supervised to all the experiments performed in this work. PT and VG performed the metabolomic experiments. BC and LC performed preliminary transcriptomic experiments. MG contributed during growth and metabolomic experiments and RNAseq data analysis under the supervision of EP and MF. CF contributed to the realization of growth experiments. MF performed/supervised to all the computation reported in this work. MF and FDP wrote the theoretical kinetic models and performed the simulations. MF wrote the manuscript. All the authors contributed to the editing of the manuscript. All the authors have read and approved the final version of the manuscript.

## Notes

#### Summary of Updates

Typos and figure numbering fixed

