## Supplementary Material S1 for "Diauxie and co-utilization are not exclusive during growth in nutritionally complex environments"

**RNA sequencing details**

The 20 samples obtained were sequenced at Genomix4life (Naples, Italy, www.genomix4life.com). The main features of the sequencing run are outlined in Table S1 below.

**Table S1.** Main features of the sequencing run obtained in this work.

| *Time point* | ***Replicate*** | ***N. of reads*** | ***NCBI accession code*** | ***File name*** |
|---|---|---|---|---|
| ***T2*** | 1 | 45589444 | SAMN12207305 | 1_S14_R1_001.fastq.gz |
| ***T2*** | 2 | 28516257 | SAMN12207306 | 2_S1_R1_001.fastq.gz |
| ***T2*** | 3 | 20484151 | SAMN12207307 | 3_S2_R1_001.fastq.gz |
| ***T2*** | 4 | 26905601 | SAMN12207308 | 4_S3_R1_001.fastq.gz |
| ***T4*** | 1 | 31632871 | SAMN12207309 | 5_S4_R1_001.fastq.gz |
| ***T4*** | 2 | 46766051 | SAMN12207310 | 6_S15_R1_001.fastq.gz |
| ***T4*** | 3 | 44460436 | SAMN12207311 | 7_S16_R1_001.fastq.gz |
| ***T4*** | 4 | 43330805 | SAMN12207312 | 8_S5_R1_001.fastq.gz |
| ***T5*** | 1 | 31713940 | SAMN12207313 | 9_S6_R1_001.fastq.gz |
| ***T5*** | 2 | 46981096 | SAMN12207314 | 10_S17_R1_001.fastq.gz |
| ***T5*** | 3 | 45117020 | SAMN12207315 | 11_S18_R1_001.fastq.gz |
| ***T5*** | 4 | 53527022 | SAMN12207316 | 12_S19_R1_001.fastq.gz |
| ***T6*** | 1 | 32100196 | SAMN12207317 | 17_S10_R1_001.fastq.gz |
| ***T6*** | 2 | 32739316 | SAMN12207318 | 18_S11_R1_001.fastq.gz |
| ***T6*** | 3 | 28202275 | SAMN12207319 | 19_S12_R1_001.fastq.gz |
| ***T6*** | 4 | 34848944 | SAMN12207320 | 20_S13_R1_001.fastq.gz |
| ***T7*** | 1 | 52496534 | SAMN12207321 | 13_S20_R1_001.fastq.gz |
| ***T7*** | 2 | 45537877 | SAMN12207322 | 14_S7_R1_001.fastq.gz |
| ***T7*** | 3 | 36699228 | SAMN12207323 | 15_S8_R1_001.fastq.gz |
| ***T7*** | 4 | 25020206 | SAMN12207324 | 16_S9_R1_001.fastq.gz |

**Comparison with *E. coli* starvation regulators**

Over-expressed TFs included a controller of outer membrane gene expression (OmpR), curli production activator (CsgD) and a putative siderophore transport system permease protein (YfhA). Among down-regulated genes (Supplementary Material S1, Table S3) we found the gene coding for stringent starvation protein response (*sspA*). In *E. coli* this protein is involved in resistance during prolonged starvation (including amino acid starvation) as a positive regulator of many stress-related gene. Its expression is induced during stationary phase as well as upon starvation for carbon, amino acid, nitrogen and phosphate. Additionally, *sspA* is positively regulated by *relA* [1]. In our experiment, *relA* did not change its expression significantly. Together with *relA*, *spoT* is also responsible for the accumulation of cellular ppGpp (promoted by carbon source starvation but not by amino acid starvation). Even in this case, we did not notice a significant change in *spot* expression during our experiment. Besides the afore mentioned RpoS, up-regulated TFs included an OmpR member (PSHAa0628), putatively annotated as a response regulator consisting of a CheY-like receiver domain and a winged-helix DNA-binding domain. This sequence shares significant similarity (E-value 7e-41) with proteobacterial (i.e. *E. coli*) members of the two-component regulatory system CreC/CreB involved in catabolic regulation. PSHAa0622 encodes the response regulator GlrR, belonging to the GntR regulator family and up-regulating the transcription of the *glmY* sRNA when cells enter the stationary growth phase. Finally, the AraC transcriptional regulator PSHAa1588 is the fifth most up-regulated TF

following T1-T3 transition, but no reliable functional annotation could be retrieved for this gene.
As for down-regulated TFs, *iscR* (PSHAa2672) is the one with the highest fold-change (-1.91). IscR is a transcriptional repressor of the *iscRSUA* operon, involved in assembly of Fe-S clusters. According to RegPrecise database, it is predicted to regulate the expression of 11 genes. PSHAa2995 (log2FC = -1.47), the forth most down-regulated gene putatively encodes a GlpR homolog, a transcriptional regulators of sugar metabolism. The third most down-regulated TF resulted to be PSHAa0390, encoding *pdhR*, the repressor of the pyruvate dehydrogenase complex. According to RegPrecise database, it is predicted to regulate the expression of 12 genes, all of them involved, at different stages in pyruvate metabolism. In *E. coli* the decreased pyruvate concentration means a decreased inhibition of PdhR [2]. While PdhR represses the transcription of its target genes, the pyruvate-bound state of the regulator is not able to bind DNA. PdhR controls (among the others) the transcription of the multi-enzyme complex of the pyruvate dehydrogenase complex [3]. Only one TF resulted to be differentially expressed (up-regulated) following the second lag phase, PSHAa1181, the sigma factor PSHAa0691.

Together with TFs, two-component regulatory systems (TCRSs) are a basic stimulus-response coupling mechanism to sense and react to changes in environmental conditions, e.g. nutrient concentration. For this reason, we identified differentially expressed TCRSs in the two selected contrasts. Overall, we found 21 TCRSs-related genes that were differentially expressed in T1 vs. T2. PSHAa0620 was the TCRS displaying the higher $\log_2$FC (1.9) following T1 to T2 transition, encoding a signal transduction histidine kinase belonging to the BaeS family, putatively involved in cell envelope stress response. PSHAa0134 (1.19 $\log_2$FC following T1-T2 transition) belongs to the VieA TCRS family and embeds both a REC and an EAL domain. The first is typical of two-component signal transduction systems enabling bacteria to sense, respond, and adapt to a wide range of environments, stressors, and growth conditions, whereas the second might be involved in regulating cell surface adhesiveness in bacteria [4].
Among the other over-expressed TCRS, we identified PSHAa2620, encoding a PleD-like response regulator, a two-component response regulator embedding two REC domains and a diguanylate cyclase (GGDEF) domain. Members of this family are the best-characterized regulator of c-di-GMP levels and motility in *C. crescentus* and have similarly been shown to be involved in regulating surface motility in other bacterial species [5–8]. PSHAa0628 (the fourth most expressed TCRS) encodes a proteobacterial dedicated sortase system response regulator (*pdsR*). This family of DNA-binding response regulator proteins are usually associated with an adjacent histidine kinase to form a two-component system; PSHAa0628 makes no exception as it is flanked by PSHAa0629, a signal transduction histidine kinase (COG0642). The other up-regulated TCRS representatives included a nitrate/nitrite response regulator NarL (PSHAa2948), an osmolarity sensing histidine kinase (PSHAa2849), a CheY-like regulator (PSHAa1501) and the copper resistance/phosphate regulon response regulator CusR (PSHAb0012).
The most down-regulated TCRS member identified was PSHAb0361 an already characterized C4-dicarboxylate sensor kinase, probably involved in regulating the expression of a C4-dicarboxylate transporter system comprising PSHAb0363 and PSHAb0364 [9].

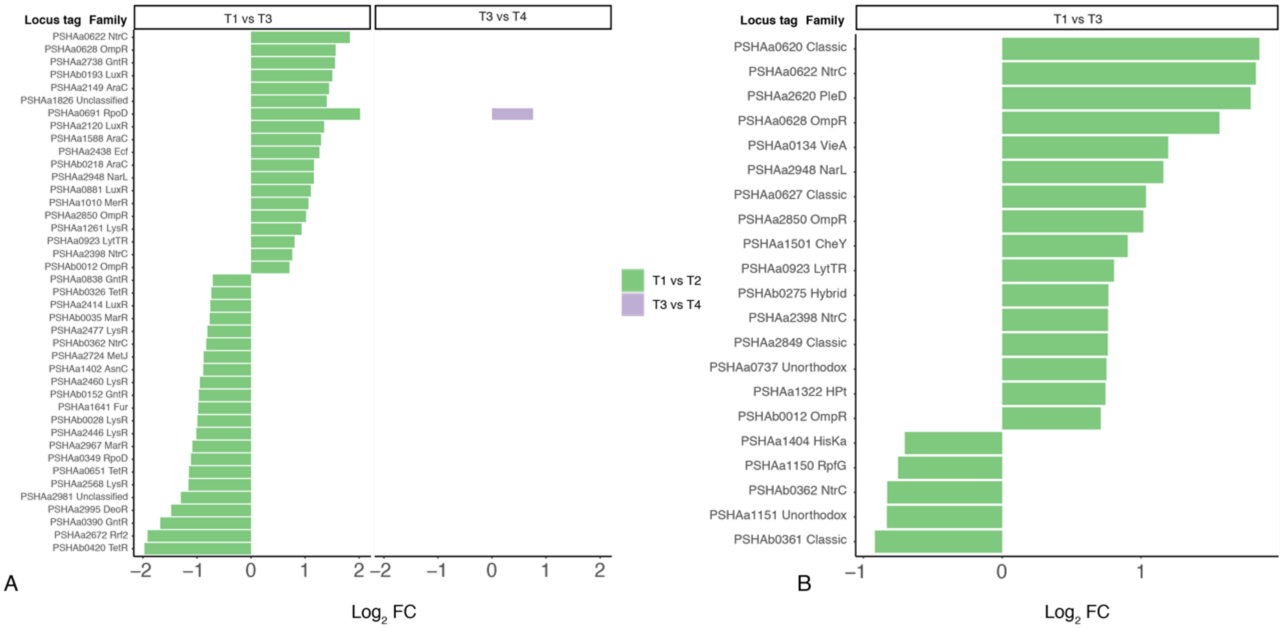


**Figure S1.** Differentially expressed transcription factors (A) and two-component regulation systems (B) in PhTAC125 following T1-T3 and T3-T5 transitions.

**Table S2.** *E. coli* global regulators known to be involved in metabolism control in *E. coli* and their corresponding PhTAC125 ortholog.

| Gene_name | Function | Gene |
|---|---|---|
| ArcB | sensor of redox conditions, activates ArcA | PSHAa0550 |
| ArgR | repressor of arginine biosynthetic genes | PSHAa2659 |
| AscG_CytR | regulator of the cryptic asc operon for β-glucoside utilization | PSHAa1771 |
| BasR | regulator of two component system; putative role is lipopolysaccharides regulation | PSHAb0162 |
| BolA | regulator of cell wall biosynthetic enzymes with different roles in cell morphology and cell division | PSHAa2243 |
| CpxR | regulator of two component system that responds to envelope stress | PSHAa0375 |
| Cra | cAMP independent regulation of catabolite-sensitive operons | PSHAa1365 |
| CreB | catabolic regulation | PSHAa2195 |
| CsgD | activator of the csgBA and csgDEFG operons necessary for curli production | PSHAa2120 |
| CspA | cold shock protein; transcriptional activator of hns | PSHAb0078 |
| CysB | regulator of cystein regulon (sulphur utilization and sulphonate-sulphur metabolism) | PSHAa1162 |
| FadR | regulator of fatty acid metabolism | PSHAa0838 |
| FhlA | FhlA is the transcriptional activator that directly controls the hycABCDEFGHI operon in the absence of molybdate, which is involved in the induction of formate hydrogen-lyase | PSHAa1893 |

| | | |
|---|---|---|
| Fis | regulator of stable RNA (rRNA and tRNA) synthesis | PSHAa0346 |
| Fnr | regulator of proteins involved in cellular adaptation to growth in anoxic environments | PSHAa1850 |
| Fur | regulator of ferric uptake | PSHAa1641 |
| GcvA | regulator of gcvTHP operon, wich is supposidely involved in balancing the requirements of the cell for glycine and C1 units | PSHAa1064 |
| HdfR | negatively regulates the expression of the flagellar master operon, flhDC A direct activator of the flhDC operon and represses transcription of hdfR | PSHAa2568 |
| IdnR | regulator of gluconate and L-idonate metabolism | PSHAb0478 |
| IHF A | integration host factor α subunit, histone-like protein | PSHAa1902 |
| IHF B | integration host factor β subunit, histone-like protein | PSHAa1426 |
| LldR/GlcC/PdhR | Lactate regulator A transcription factor/ GlcC-Glycolate transcriptional dual regulator/ pyruvate dehydrogenase complex regulator | PSHAa0390 |
| Lrp | leucine responsive regulatory protein; regulator of high-affinity branched –chain amino acid transport system | PSHAa1717 |
| NsrR | NsrR is a transcriptional regulator that negatively regulates the transcription of ytfE, hmpA, and ygbA Expression of these genes is normally upregulated by nitrosative stress | PSHAa2419 |
| NtrC | regulator of genes involved in nitrogen assimilation | PSHAa0163 |
| OmpR | regulator participates in controlling the expression of major outer membrane genes | PSHAa2850 |
| OxyR | regulator that belongs to the LysR family and participates in controlling several genes involved in the response to oxidative stress and the production of surface proteins that control the colony morphology and auto-aggregation ability | PSHAa0760 |
| PhoB | regulation of the phosphate regulon | PSHAa0597 |
| QseB | putative 2-component transcriptional regulator involved in the regulation of flagella and motility by quorum sensing | PSHAa1283 |
| SspA | stringent starvation protein | PSHAa2527 |
| TorR | regulator of two component system involved in trimethylamine N-oxide (TMAO) anaerobic respiration | PSHAa0551 |
| UvrY | regulator of two component system regulation genes involved in DNA repair and carbon metabolism | PSHAa1916 |
| YdjF | putative transcriptional regulator | PSHAb0550 |
| YfhA | putative two component transcriptional regulator | PSHAa2620 |

**Table S3.** Differentially expressed global metabolic regulators in *Ph*TAC125.

| **Locus tag** | **Product name** | **Function** | **$Log_{10}$ FC** | **Adj. P-value** |
|---|---|---|---|---|
| PSHAa2850 | OmpR | regulator participates in controlling the expression of major outer membrane genes | 1.01 | 8.37e-22 |

| | | | | |
|---|---|---|---|---|
| PSHAa2120 | CsgD | activator of the csgBA and csgDEFG operons necessary for curli production | 1.34 | 1.03e-11 |
| PSHAa0622 | YfhA | putative two component transcriptional regulator | 1.82 | 3.68e-66 |
| PSHAa0346 | Fis | regulator of stable RNA (rRNA and tRNA) synthesis | -2.17 | 2.86e-45 |
| PSHAa0390 | LldR_GlcC_PdhR | Lactate regulator A transcription factor that controls expression of genes involved in transport and catabolism of L-lactate | -1.67 | 9.88e-72 |
| PSHAb0078 | CspA | cold shock protein; transcriptional activator of hns | -1.34 | 1.21e-11 |
| PSHAa2568 | HdfR | negatively regulates the expression of the flagellar master operon, flhDC A direct activator of the flhDC operon and represses transcription of hdfR | -1.15 | 5.77e-23 |
| PSHAa2527 | SspA | stringent starvation protein | -0.98 | 2.12e-31 |
| PSHAa1641 | Fur | regulator of ferric uptake | -0.97 | 2.39e-11 |
| **PSHAa0838** | FadR | regulator of fatty acid metabolism | -0.70 | 3.00e-12 |

**Table S4.** Global transcriptional regulators in *E. coli*, their ortholog in *Ph*TAC125 and whether they were differentially expressed or not in our experiment.

| ***E. coli* regulator** | ***E. coli* locus tag** | ***Ph*TAC125 closest homolog** | **DEG in *Ph*TAC125 (T1 vs. T2/T3 vs. T4)** |
|---|---|---|---|
| *mlc* | b1594 | PSHAb0149 | No/No |
| *rpoS (sigma 38)* | b2741 | PSHAa0691 | Yes/Yes |
| *FecI (sigma 19)* | | - | |
| *rpoE (sigma 24)* | b2573 | PSHAa0726 | No/No |
| *fliA (sigma 38)* | b1922 | PSHAa0809 | No/No |
| *rpoH (sigma 32)* | b3461 | PSHAa0357 | No/No |
| *rpoN (sigma 54)* | b3202 | PSHAa2551 | No/No |
| *rpoD (sigma 70)* | b3067 | PSHAa0349 | Yes/No |

**Table S5.** List of differentially expressed *Ph*TAC125 TFs identified. All differentially expressed TS were identified following the T1-T3 transition, except * that was identified following the T3-T5 one

| Locus Tag | TF family | Log2 FC | Adj. P-value | General description |
|---|---|---|---|---|
| PSHAa0691 | RpoD | 2.01 | 8.15e-70 | Sigma factor, RpoD family contains 1 Sigma70_r1_2,1 Sigma70_r2,1 Sigma70_r3,1 Sigma70_r4 |
| PSHAa0622 | NtrC | 1.82 | 3.68e-66 | Response regulator, NtrC family contains 1 Response_reg,1 AAA,1 HTH_8 |
| PSHAa0628 | OmpR | 1.56 | 1.58e-34 | Response regulator, OmpR family contains 1 Response_reg,1 Trans_reg_C |
| PSHAa2738 | GntR | 1.55 | 5.58e-10 | Transcription factor, GntR family contains 1 GntR,1 UTRA |
| PSHAb0193 | LuxR | 1.50 | 1.73e-31 | Transcription factor, LuxR family contains 1 HTH_LUXR |
| PSHAa2149 | AraC | 1.44 | 1.96e-13 | Transcription factor, AraC family contains 1 HTH_AraC |
| PSHAa1826 | Unclassified | 1.39 | 1.60e-28 | Sigma factor, Unclassified contains 1 Sigma70_r2,1 HTH_LUXR |
| PSHAa2120 | LuxR | 1.34 | 1.03e-11 | Transcription factor, LuxR family contains 1 HTH_LUXR |
| PSHAa1588 | AraC | 1.29 | 3.04e-30 | Transcription factor, AraC family contains 1 AraC_N,2 HTH_AraC |
| PSHAa2438 | Ecf | 1.26 | 3.86e-07 | Sigma factor, Ecf family contains 1 Sigma70_r2,1 Sigma70_r4 |
| PSHAb0218 | AraC | 1.16 | 1.17e-17 | Transcription factor, AraC family contains 1 AraC_N,1 HTH_AraC |
| PSHAa2948 | NarL | 1.16 | 4.49e-22 | Response regulator, NarL family contains 1 Response_reg,1 HTH_LUXR |
| PSHAa0881 | LuxR | 1.10 | 8.19e-18 | Transcription factor, LuxR family contains 1 HTH_LUXR |
| PSHAa1010 | MerR | 1.06 | 1.53e-10 | Transcription factor, MerR family contains 1 MerR,1 MerR-DNA-bind |
| PSHAa2850 | OmpR | 1.01 | 8.37e-22 | Response regulator, OmpR family contains 1 Response_reg,1 Trans_reg_C |
| PSHAa1261 | LysR | 0.93 | 3.41e-07 | Transcription factor, LysR family contains 1 HTH_1,1 LysR_substrate |
| PSHAa0923 | LytTR | 0.80 | 1.19e-05 | Response regulator, LytTR family contains 1 Response_reg,1 LytTR |
| PSHAa2398 | NtrC | 0.76 | 2.53e-15 | Response regulator, NtrC family contains 1 Response_reg,1 AAA_5,1 HTH_8 |

| | | | | |
|---|---|---|---|---|
| PSHAb0012 | OmpR | 0.70 | 0.0003 | Response regulator, OmpR family contains 1 Response_reg,1 Trans_reg_C |
| PSHAa0838 | GntR | -0.70 | 3.00e-12 | Transcription factor, GntR family contains 1 GntR |
| PSHAb0326 | TetR | -0.73 | 9.14e-08 | Transcription factor, TetR family contains 1 TetR_N |
| PSHAa2414 | LuxR | -0.75 | 1.097e-05 | Transcription factor, LuxR family contains 1 HTH_LUXR |
| PSHAb0035 | MarR | -0.76 | 9.31e-08 | Transcription factor, MarR family contains 1 MarR |
| PSHAa2477 | LysR | -0.80 | 1.04e-06 | Transcription factor, LysR family contains 1 HTH_1,1 LysR_substrate |
| PSHAb0362 | NtrC | -0.82 | 6.19e-14 | Response regulator, NtrC family contains 1 Response_reg,1 AAA_5,1 HTH_8 |
| PSHAa2724 | MetJ | -0.87 | 6.80e-17 | Transcription factor, MetJ family contains 1 MetJ |
| PSHAa1402 | AsnC | -0.88 | 3.62e-14 | Transcription factor, AsnC family contains 1 Rrf2,1 AsnC_trans_reg |
| PSHAa2460 | LysR | -0.94 | 2.97e-16 | Transcription factor, LysR family contains 1 HTH_1,1 LysR_substrate |
| PSHAb0152 | GntR | -0.96 | 1.02e-18 | Transcription factor, GntR family contains 1 GntR |
| PSHAa1641 | Fur | -0.97 | 2.39e-11 | Transcription factor, Fur family contains 1 FUR |
| PSHAb0028 | LysR | -0.99 | 2.18e-15 | Transcription factor, LysR family contains 1 HTH_1,1 LysR_substrate |
| PSHAa2446 | LysR | -1.01 | 2.23e-11 | Transcription factor, LysR family contains 1 HTH_1,1 LysR_substrate |
| PSHAa2967 | MarR | -1.08 | 1.35e-35 | Transcription factor, MarR family contains 1 MarR |
| PSHAa0349 | RpoD | -1.11 | 3.49e-41 | Sigma factor, RpoD family contains 1 Sigma70_r1_1,1 Sigma70_r1_2,1 Sigma70_ner,1 Sigma70_r2,1 Sigma70_r3,1 Sigma70_r4 |
| PSHAa0651 | TetR | -1.14 | 1.88e-17 | Transcription factor, TetR family contains 1 TetR_N |
| PSHAa2568 | LysR | -1.15 | 5.77e-23 | Transcription factor, LysR family contains 1 HTH_1,1 LysR_substrate |
| PSHAa2981 | Unclassified | -1.29 | 1.15e-09 | Transcription factor, Unclassified contains 1 HTH_12 |
| PSHAa2995 | DeoR | -1.47 | 3.75e-20 | Transcription factor, DeoR family contains 1 HTH_DeoR,1 DeoR |
| PSHAa0390 | GntR | -1.67 | 9.88e-72 | Transcription factor, GntR family contains 1 GntR |
| PSHAa2672 | Rrf2 | -1.91 | 1.69e-63 | Transcription factor, Rrf2 family contains 1 Rrf2 |

| | | | | |
|---|---|---|---|---|
| PSHAb0420 | TetR | -1.97 | 2.09e-27 | Transcription factor, TetR family contains 1 TetR_N |
| PSHAa0691* | RpoD | 0.75 | 2.74e-28 | Sigma factor, RpoD family contains 1 Sigma70_r1_2,1 Sigma70_r2,1 Sigma70_r3,1 Sigma70_r4 |

**Table S6.** List of differentially expressed *Ph*TAC125 TCSRs identified. All differentially expressed TS were identified following the T1-T3 transition, except * that was also identified following the T3-T5 one ($log_2$FC and p-value are provided in the same line)

| Locus Tag | TCRS family | $Log_2$ FC (T1-T2/T3-T4) | Adj. P-value | General description |
|---|---|---|---|---|
| PSHAa0620 | Classic | 1.85 | 5.54e-85 | Histidine kinase, Classic contains 1 HAMP,1 HisKA,1 HATPase_c |
| PSHAa0622 | NtrC | 1.82 | 3.68e-66 | Response regulator, NtrC family contains 1 Response_reg,1 AAA,1 HTH_8 |
| PSHAa2620 | PleD | 1.78 | 3.95e-90 | Response regulator, PleD family contains 1 Response_reg,1 GGDEF |
| PSHAa0628 | OmpR | 1.56 | 1.58e-34 | Response regulator, OmpR family contains 1 Response_reg,1 Trans_reg_C |
| PSHAa0134 | VieA | 1.19 | 5.57e-17 | Response regulator, VieA family contains 1 Response_reg,1 EAL |
| PSHAa2948 | NarL | 1.16 | 4.49e-22 | Response regulator, NarL family contains 1 Response_reg,1 HTH_LUXR |
| PSHAa0627 | Classic | 1.03 | 2.12e-09 | Histidine kinase, Classic contains 1 HAMP,1 HisKA,1 HATPase_c |
| PSHAa2850 | OmpR | 1.01 | 8.37e-22 | Response regulator, OmpR family contains 1 Response_reg,1 Trans_reg_C |
| PSHAa1501 | CheY | 0.90 | 4.28e-05 | Response regulator, CheY family contains 1 Response_reg |
| PSHAa0923 | LytTR | 0.80 | 1.19e-05 | Response regulator, LytTR family contains 1 Response_reg,1 LytTR |
| PSHAb0275 | Hybrid | 0.76 | 1.53e-07 | Histidine kinase, Hybrid contains 1 HisKA,1 HATPase_c,1 Response_reg |
| PSHAa2398 | NtrC | 0.76 | 2.53e-15 | Response regulator, NtrC family contains 1 |

| | | | | |
|---|---|---|---|---|
| | | | | Response_reg,1 AAA_5,1 HTH_8 |
| PSHAa2849 | Classic | 0.76 | 1.74e-08 | Histidine kinase, Classic contains 1 HAMP,1 HisKA,1 HATPase_c |
| PSHAa0737 | Unorthodox | 0.75 | 7.99e-07 | Histidine kinase, Unorthodox contains 1 HAMP,1 HisKA,1 HATPase_c,1 Response_reg,1 Hpt |
| PSHAa1322 | HPt | 0.74 | 5.6e-08 | Phosphotransfer protein contains 1 Hpt |
| PSHAb0012 | OmpR | 0.70 | 0.0003 | Response regulator, OmpR family contains 1 Response_reg,1 Trans_reg_C |
| PSHAa1404 | HisKa | -0.70 | 1.02e-14 | Phosphotransfer protein contains 1 His_kinase |
| PSHAa1150 | RpfG | -0.74 | 6.24e-12 | Response regulator, RpfG family contains 1 Response_reg,1 HD |
| PSHAb0362 | NtrC | -0.82 | 6.19e-14 | Response regulator, NtrC family contains 1 Response_reg,1 AAA_5,1 HTH_8 |
| PSHAa1151 | Unorthodox | -0.83 | 3.70-12 | Histidine kinase, Unorthodox contains 3 MHYT,1 PAS_3,1 HisKA,1 HATPase_c,1 Response_reg,1 Hpt |
| PSHAb0361 | Classic | -0.91 | 3.54e-18 | Histidine kinase, Classic contains 1 HisKA,1 HATPase_c |

**Main players of amino acid metabolism in a complex medium**

The set of amino acids that are typically consumed in the first ours of PhTAC125 growth comprises Ser, Thr, Asp, Asn and Glu. Afterwards, PhTAC125 metabolism switches to the consumption of a second set of amino acids that includes Lys, Leu, Ala, Gly, Phe, Tyr, Ile and Val. In the experimental set up described by Wilmes et al. (2010), this switch occurs after 4 hours. Four additional hours are then required to consume this second set of amino acids, and to finally start degrading the amino acid His. No information is currently available on the faith of Met, Cys, Trp, Pro, Gln and Arg during growth of PhTAC125 in a complex medium.

Indeed, Ser, Thr, Asn, Asp and Glu were shown to be the first amino acids to be consumed in an amino acid rich medium; consistently, we found the down-regulation of *purA*, *murI*, and *sdaA* involved in the degradation of Asp, Glu and Ser, respectively. Interestingly, 4 genes out of 7 of the down-regulated and 4 out of 6 of the up-regulated between T1 and T3 are involved in the first utilization step of the same amino acids, i.e. Asp, Glu, Met and Ser (Figure 2E and F). Considering the genes analysed here, Asp seems to be converted to andenylosuccinate at T1, then this gene is turned off and the degradation redirected towards the production of Arg. Similarly, the gene converting Ser to pyruvate (*sdaA*, L-serine ammonia-lyase) is predicted to be significantly down-regulated following T1 to T3 transition, whereas the genes involved in the conversion of Ser to Gly (*glyA*) shows a significant increase in the same contrast. Glutamate racemase *(murI)*, responsible for the conversion of L-Glu to D-Glu appears to be turned off between T1 and T3, whereas the expression of the gene responsible for the conversion of Glu to 2-oxo-glutarate (*gdhA*) shows a significant increase. Finally, *mdeA* and *metK* are significantly up- and dow-regulated, respectively, when considering the T1 vs T3 time interval. The first is responsible for the conversion of Met to methanediol, allowing the entrance of this intermediate into sulfur metabolism, whereas the second encodes the conversion of Met to S-adenosyl-methionine (SAM).

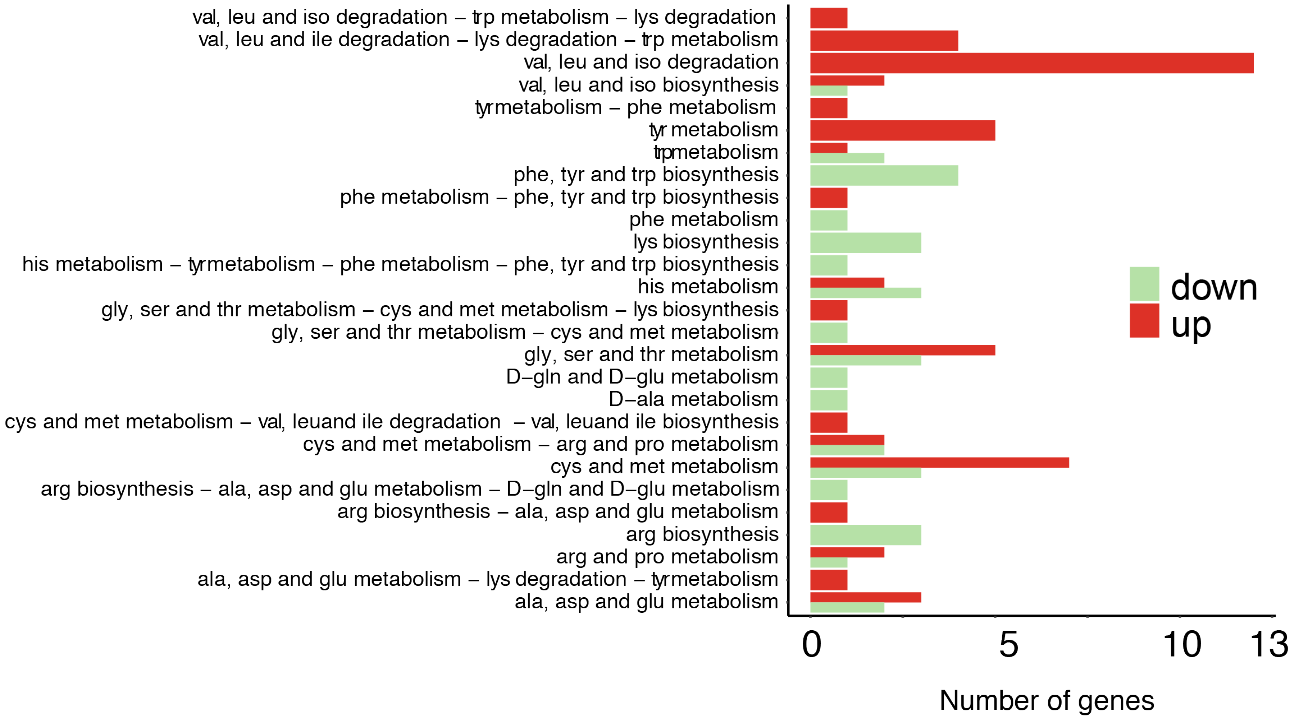


**Figure S2.** Number of differentially expressed amino acid metabolic genes for each pathway

**Table S7.** List of genes responsible for the first assimilation step of amino acids in *Ph*TC125.

| **Locus Tag and function** | **Gene name** | **Assimilation pathway** |
|---|---|---|
| PSHAa1163 sdaA L-serine deaminase I | *sdaA* | Ser |
| PSHAa2768 ilvA, threonine dehydratase | *ilvA* | Thr |
| PSHAa1288 trpA, tryptophan synthase, alpha protein | *trpA* | Ser |
| PSHAa1289 trpB, tryptophan synthase, beta protein | *trpB* | Ser |
| PSHAa2315 threonine 3-dehydrogenase | *tdh* | Thr |
| PSHAa1408 L-asparaginase IL-asparagine amidohydrolase I | *ansA* | Asp |
| PSHAa1440 asparagine synthetase Bglutamine-hydrolyzing | *asnB* | Asp/Asn |
| PSHAa2796 asparagine synthetase A | *asnA* | Asp |

| | | |
|---|---|---|
| PSHAa0725 quinolinate synthetase B protein | *nadB* | Asp |
| PSHAa0956 aspartate aminotransferase Transaminase A | *aspC* | Asp/Thr |
| PSHAa2288 Argininosuccinate synthase Citrulline--aspartate ligase | *argG* | Asp |
| PSHAa0275 adenylosuccinate synthetase | *purA* | Asp |
| PSHAa1692 adenylosuccinate lyase | *purB* | Asp |
| PSHAb0019 putative dinucleotide-utilizing enzyme | *PSHAb0019* | Asp |
| PSHAa0533 Aspartokinase | *lysC* | Asp |
| PSHAa1250 lysine-sensitive aspartokinase III | *lysC2* | Asp |
| PSHAa2379 Bifunctional aspartokinase/homoserine dehydrogenase I | *thrA* | Asp |
| PSHAa2722 bifunctional: aspartokinase II homoserine dehydrogenase II | *metL* | Asp |
| PSHAa2293 putative cysteine sulfinic acid decarboxylase | *PSHAa2293* | Glu |
| PSHAa0166 glutamine synthetase | *glnA* | Glu |
| PSHAa2264 Proline dehydrogenase | *putA* | Glu/Pro |
| PSHAa1670 putative glutamate dehydrogenase | *PSHAa1670* | Glu |
| PSHAa1392 glutamate dehydrogenase, NADP-specific | *gdhA* | Glu |
| PSHAa0252 glutamate racemase | *murI* | Glu |
| PSHAa0937 glutamate--cysteine ligase Gamma-glutamylcysteine synthetase | *gshA* | Glu |
| PSHAa1094 putative basic aminoacid decarboxylase | *PSHAa1094* | Lys |
| PSHAa1167 Leucine dehydrogenase | *bcd* | Leu |
| PSHAa1270 putative branched-chain amino acid aminotransferase | *ilvE* | Ile/Val/Leu |
| PSHAa1323 putative PLP-dependent aminotransferase | *yfbQ* | Ala |
| PSHAa2434 alanine racemase | *alr* | Ala |
| PSHAa2473 glycine cleavage complex protein P, glycine decarboxylase, PLP-dependent | *gcvP* | Gly |
| PSHAb0295 Low-specificity L-threonine aldolase | *ltaE* | Gly/Ser |
| PSHAa2376 glyA, serine hydroxymethyltransferase | *glyA* | Ser |
| PSHAa2043 Phenylalanine-4-hydroxylase EC 1.14.16.1 | *phhA* | Phe |
| PSHAb0492 histidinol phosphate aminotransferase | *hisC* | Thr |
| PSHAa2740 Histidine ammonia-lyase | *hutH* | His |
| PSHAa0670 S-adenosylmethionine synthetase Met | *metK* | Met |
| PSHAa2044 methionine-gamma-lyase Met | *mdeA* | Met |
| PSHAa2605 pyrroline-5-carboxylate reductase Pro | *proC* | Pro |
| PSHAa2846 arginine decarboxylase Arg | *speA* | Arg |
| PSHAa0195 arginine N-succinyltransferase | *astA* | Arg |
| PSHAa1075 arginine decarboxylase | *speA* | Arg |

**Amino acids metabolic pathways regulation**

The expression of all the genes involved in AA metabolism in PhTAC125 was evaluated (Figure S3). Each gene was associated to a specific amino acid metabolic pathway according to the KEGG database and the correlation (Pearson correlation) among each gene belonging to the same pathway was evaluated (Table S8). In most cases, we found a low level of (positive) correlation among the expression values of the genes belonging the same metabolic pathway. Most of the Fisher's z-transformed correlation coefficient range between 0.5 and 0.14. Genes involved in Tyr metabolism and branched chain amino acid degradation are the ones that display the highest correlation values, with 1.15 and 1.07. respectively. Conversely, Phe, Tyr and Trp biosynthetic genes were those with the lowest correlation (0.14).

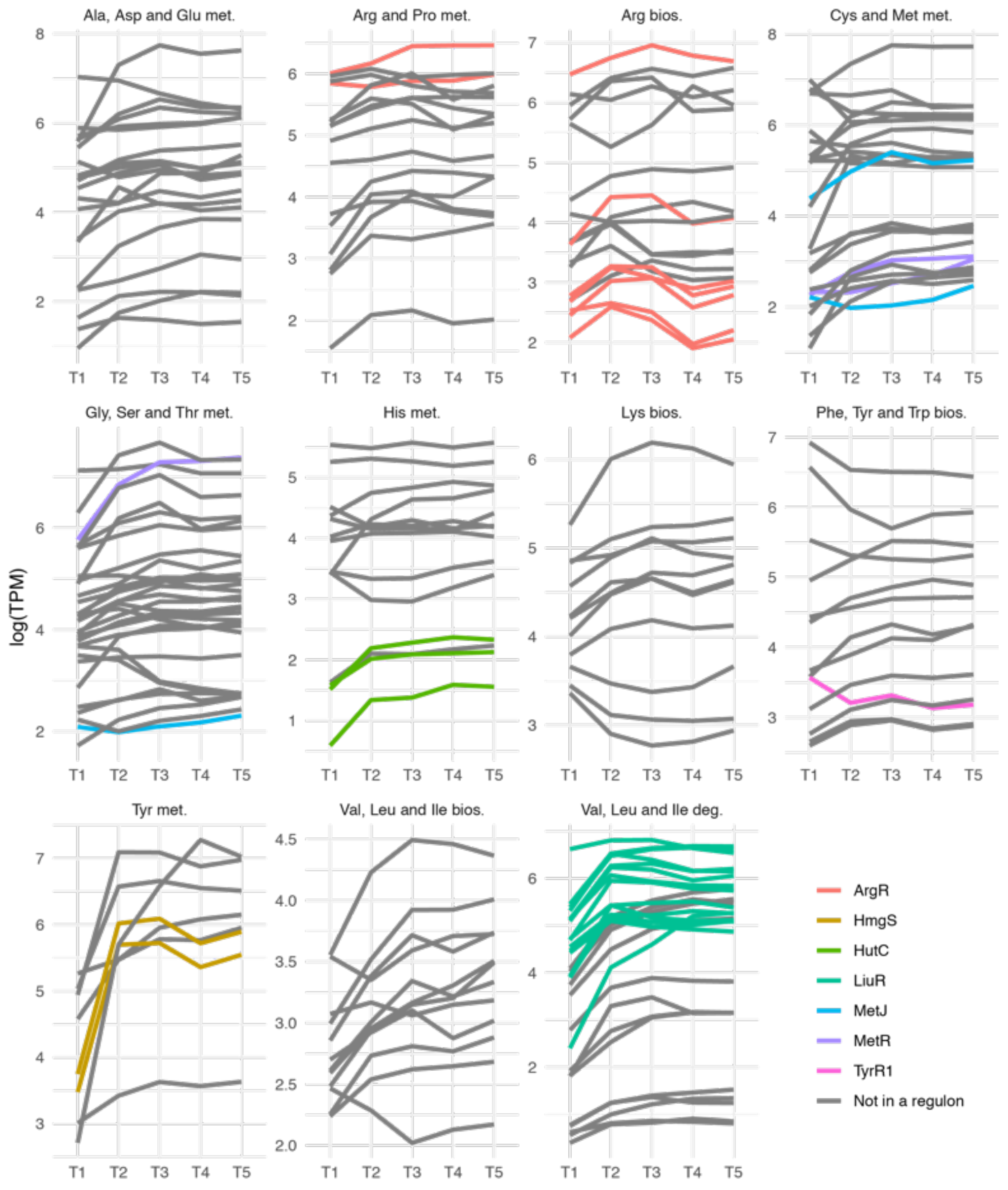


**Figure S3.** Gene expression pattern of all the genes involved in amino acid metabolism in *Ph*TAC125. Coloured lines represent genes assigned to a specific regulon (see legend) according to the RegPrecise database.

**Table S8.** Fisher's Z transformation average of Pearson correlation coefficient among the genes belonging to the same amino acid metabolic pathway.

| **Amino acid metabolic pathway** | **Fisher's z transformation of Pearson correlation coefficient** |
|---|---|
| Tyr met. | 1,15 |
| Val, Leu and Ile deg. | 1,07 |
| Val, Leu and Ile bios. | 0,59 |
| Arg and Pro met. | 0,511 |
| Ala, Asp and Glu met. | 0,51 |
| Gly, Ser and Thr met. | 0,44 |
| Cys and Met met. | 0,29 |
| Arg bios. | 0,271 |
| His met. | 0,23 |
| Lys bios. | 0,23 |
| Phe, Tyr and Trp bios. | 0,14 |

**Table S9.** Main information for all the *Ph*TAC125 regulons considered in this work.

| Regulator | Effector | Pathway | Operons |
|---|---|---|---|
| ArgR | Arginine | Arginine biosynthesis; Arginine degradation | 4 |
| HmgS | | Tyrosine degradation | 2 |
| HutC | cis-Urocanic acid | Histidine utilization | 2 |
| LiuR | | Branched-chain amino acid degradation | 9 |
| MetJ | S-adenosylmethionine | Methionine metabolism; Methionine biosynthesis | 9 |
| MetR | Homocysteine | Methionine biosynthesis | 4 |
| TyrR1 | Tyrosine; Phenylalanine | Aromatic amino acid metabolism | 4 |

### Main players in amino acids assimilation

We selected one amino acid for each of the four groups (Glu, Asp, Phe and Met) and analysed the expression (delta-CT values) of those genes that i) were involved in the their possible first assimilatory steps (Figure 3D and Supplementary Material Table S2) and ii) resulted to be differentially expressed in, at least, one contrast of the differential transcriptomic analysis. Data obtained (see Supplementary file S1 for details on the analysis of RT-PCR data) revealed interesting insights on the regulation of amino acid metabolism during nutrients switching. Glutamine synthetase (*glnA*), for example, shows an increase in its expression until the first lag point (4 hours); afterwards, its expression decreases (Figure S4). GlnA uses Glu as substrate, thus this expression trend is consistent with the concentration of Glu dropping close to zero after four hours of growth. The expression increase in the first stage of the growth can be explained by the ongoing exhaustion of Gln in the medium and the necessity to synthesize it from Glu. Further, *asnB* encodes an asparagine synthetase B and is involved in the formation of Asn from Asp. The expression of this gene dramatically increases following the second short lag phase reported (i.e. after 6 hours of growth) (Figure S4). This is consistent with the exhaustion of Asn in the medium at that time point and with the necessity to synthesize this amino acid, i.e. adding an amino group to Asp that, in turn, can be synthesized from TCA intermediates (e.g. oxaloacetate). *metK* (Met degradation) displays a trend that is characterized by an initial increase of its expression, followed by a down regulation in correspondence of the first growth lag phase, and a final expression increase consistent with the necessity to actively exploit these carbon and energy sources when most of the others are depleted. Finally, we confirmed the complementary expression pattern of *metK* and *mdeA*, responsible for the entrance of Met degradation intermediates into sulfur metabolism and for the conversion of Met to S-adenosyl-methionine (SAM), respectively (Figure S4).

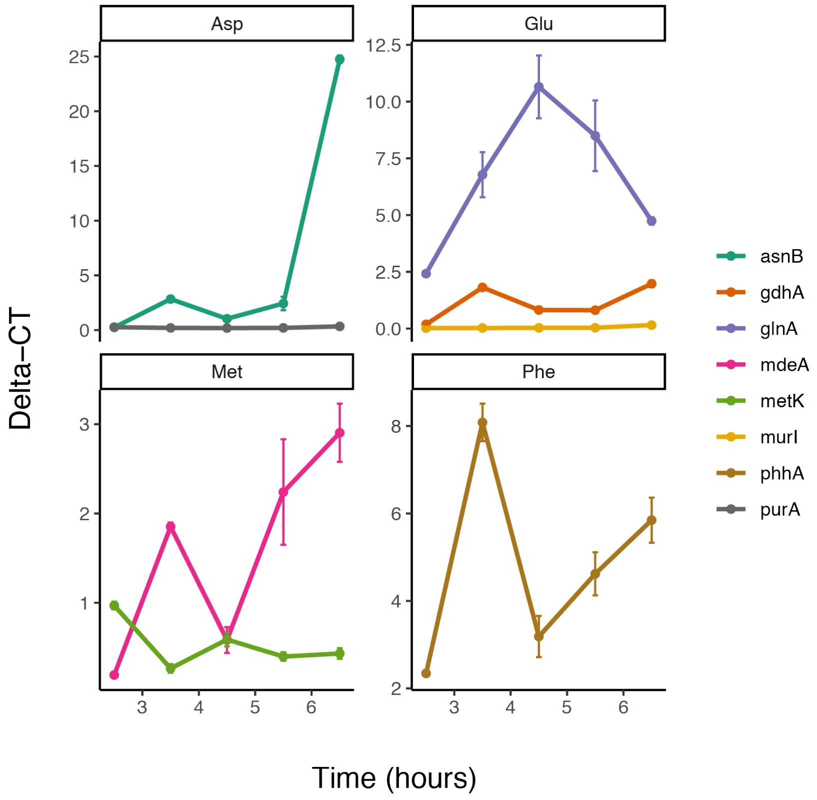


**Figure S4.** Expression (delta-CT) values for the genes involved in the first assimilatory steps of 4 selected amino acids.

### Model parameters

**Table S10.** Parameters used in this work for the MMM model

| Parameter | Description | Value | Units |
|---|---|---|---|
| $\beta_1$ | max rate constant of *P* on $S_1$ | 0.0919 | $g/g_{CDW}*h$ |
| $\beta_2$ | max rate constant of *P* on $S_2$ | 0.0478 | $g/g_{CDW}*h$ |
| $\beta_3$ | max rate constant of *P* on $S_3$ | 0.3101 | $g/g_{CDW}*h$ |
| $\beta_4$ | max rate constant of *P* on $S_4$ | 0.0277 | $g/g_{CDW}*h$ |
| $k_1$ | Michaelis-Menten constant for $S_1$ | 0.0497 | g/l |

| | | | |
|---|---|---|---|
| $k_2$ | Michaelis-Menten constant for $S_2$ | 0.0268 | g/l |
| $k_3$ | Michaelis-Menten constant for $S_3$ | 2.9170 | g/l |
| $k_4$ | Michaelis-Menten constant for $S_4$ | 3.3195 | g/l |
| $d$ | Death rate | 0.001 | $h^{-1}$ |

**Table S11.** Parameters used in this work for the cybernetic model

| Parameter | Description | Value | Units |
|---|---|---|---|
| $V_{max1}$ | max rate constant of $P$ on $S_1$ | 0.20341 | g/$g_{CDW}$*h |
| $V_{max2}$ | max rate constant of $P$ on $S_2$ | 2.3651 | g/$g_{CDW}$*h |
| $V_{max3}$ | max rate constant of $P$ on $S_3$ | 160.45 | g/$g_{CDW}$*h |
| $V_{max4}$ | max rate constant of $P$ on $S_4$ | 0.35559 | g/$g_{CDW}$*h |
| $K_{s1}$ | Michaelis-Menten constant for $S_1$ | 182.06 | g/l |
| $K_{s2}$ | Michaelis-Menten constant for $S_2$ | 5.0622 | g/l |
| $K_{s3}$ | Michaelis-Menten constant for $S_3$ | 4484.6 | g/l |
| $K_{s4}$ | Michaelis-Menten constant for $S_4$ | 39.41 | g/l |
| $V_{e1}$ | max rate constant of $e_1$ synthesis | 1.8702 | g/$g_{CDW}$*h |
| $V_{e2}$ | max rate constant of $e_2$ synthesis | 1.0252e-05 | g/$g_{CDW}$*h |
| $V_{e3}$ | max rate constant of $e_3$ synthesis | 0.020896 | g/$g_{CDW}$*h |
| $V_{e4}$ | max rate constant of $e_4$ synthesis | 0.014356 | g/$g_{CDW}$*h |
| $K_{e1}$ | Michaelis-Menten constant for $e_1$ | 0.0034958 | g/l |
| $K_{e2}$ | Michaelis-Menten constant for $e_2$ | 2.0675e-08 | g/l |
| $K_{e3}$ | Michaelis-Menten constant for $e_3$ | 8.4553e-13 | g/l |
| $K_{e4}$ | Michaelis-Menten constant for $e_4$ | 2.4577e-08 | g/l |
| $d$ | Death rate | 0.001 | $h^{-1}$ |
| $\alpha$ | Enzyme dilution rate | 0.0098085 | $h^{-1}$ |
| $\lambda$ | P fraction increase due to consumption of S | 1 | - |
| $\beta$ | Basic enzyme synthesis rate | 0.001 | $h^{-1}$ |

### RT-PCR experiments

**Table S12.** Features of the RT primers used in this work

| Primer | Sequence | Tm (°C) | Lenght (bp) | GC% | Amplicon size (bp) | Source |
|---|---|---|---|---|---|---|
| metK_TAC_for | 5'-CGTACTAGCCCAGAAGAGCA-3' | 58.90 | 20 | 55.00 | 160 | This work |
| metK_TAC_rev | 5'-GTAACCAGTTAAGCTCGCCG-3' | 59.00 | 20 | 55.00 | | This work |
| glnA_TAC_for | 5'-TCAATTGCTGGCTGGAAAGG-3' | 58.74 | 20 | 50.00 | 151 | This work |
| glnA_TAC_rev | 5'-CGCGCTCGTAACCTTGTAAT-3' | 58.73 | 20 | 50.00 | | This work |
| purA_TAC_for | 5'-GCGCACAAGGTACGTTACTT-3' | 58.86 | 20 | 50.00 | 164 | This work |
| purA_TAC_rev | 5'-GCCTGAACCAACACGTGTAG-3' | 59.13 | 20 | 55.00 | | This work |
| murI_TAC_for | 5'-ATCAAGTCCCTATACGCGCA-3' | 58.96 | 20 | 50.00 | 150 | This work |
| murI_TAC_rev | 5'-CGATCGAGTTCTTGGTGCAG-3' | 59.01 | 20 | 55.00 | | This work |
| gdhA_TAC_for | 5'-GCTCTACCTTGGGCCCATAT-3' | 58.93 | 20 | 55.00 | 151 | This work |
| gdhA_TAC_rev | 5'-GGATCAAAGTTAGCACCGCC-3' | 59.27 | 20 | 55.00 | | This work |
| phhA_TAC_for | 5'-AATTTCCGGTGGCGACTTTT-3' | 58.68 | 20 | 45.00 | 188 | This work |
| phhA_TAC_rev | 5'-CATGCGCGCTAGGTATACAC-3' | 58.94 | 20 | 55.00 | | This work |
| mdeA_TAC_for | 5'-CGGTATCGGCCTCAGTACTT-3' | 58.97 | 20 | 55.00 | 160 | This work |
| mdeA_TAC_rev | 5'-GCGCGTAATTCATCCTCGTT-3' | 59.07 | 20 | 50.00 | | This work |
| asnB_TAC_for | 5'-TGATCCCGTACAGTTGCGTA-3' | 58.82 | 20 | 50.00 | 157 | This work |
| asnB_TAC_rev | 5'-ATATAAAGGCTGTGCACCGC-3' | 58.69 | 20 | 50.00 | | This work |

| | | | | | | |
|---|---|---|---|---|---|---|
| sdaA_TAC_for | 5'-TGCTCACCACTACCACGAAT-3' | 59.03 | 20 | 50.00 | 155 | This work |
| sdaA_TAC_rev | 5'-GAAAATGCCGCCGAAATTGG-3' | 59.00 | 20 | 50.00 | | This work |
| glyA_TAC_for | 5'-AATAGCTTCGTCGCCACATG-3' | 58.71 | 20 | 50.00 | 151 | This work |
| glyA_TAC_rev | 5'-TGTAGATATGGCGCACGTTG-3' | 58.43 | 20 | 50.00 | | This work |
| rplM_TAC_for | 5'-TGATAAACTTCAAGCTGCAAAGC-3' | 58.45 | 23 | 39.13 | 152 | This work |
| rplM_TAC_rev | 5'-GAACCTGAGGCTGTTGTGC-3' | 59.05 | 19 | 57.89 | | This work |
| dnaA_TAC_for | 5'-GCTAACAAAGAGCGCTCACA-3' | 58.85 | 20 | 50.00 | 184 | This work |
| dnaA_TAC_rev | 5'-GTGTTTCAAGCTCAGGAGGC-3' | 59.12 | 20 | 55.00 | | This work |
